# Generation of anti-tumor chimeric antigen receptors incorporating T cell signaling motifs

**DOI:** 10.1101/2022.12.25.521452

**Authors:** Lakshmi Balagopalan, Taylor Moreno, Haiying Qin, Jason Yi, Katherine M. McIntire, Neriah Alvinez, Sandeep Pallikkuth, Mariah E. Lee, Hidehiro Yamane, Andy D. Tran, Philippe Youkharibache, Raul E. Cachau, Naomi Taylor, Lawrence E. Samelson

**Author notes:** Address correspondence to (L.B.), (L.E.S). These authors contributed equally to this work (Taylor Moreno, Haiying Qin).

## Abstract

Chimeric antigen receptors (CAR) T cells have been successfully used to treat lymphoma, leukemia, and multiple myeloma, but adverse effects due to cytokine secretion, CAR-T cell exhaustion, and loss of target antigen have limited their potential. Furthermore, while CARs have been designed to harness T Cell Receptor (TCR) signaling, they are significantly less sensitive than TCRs, resulting in suboptimal signaling. We have developed novel Chimeric Adapter Proteins (CAPs) that are designed to trigger signaling downstream of the TCRζ chain. CAPs are chimeric molecules that contain adapter domains in tandem with the kinase domain of ZAP70, fused to an extracellular targeting domain. We hypothesized that CAPs would be more potent than CARs because kinetic proofreading steps that define the signaling threshold and the inhibitory regulation of upstream molecules are bypassed. Indeed, second generation CAPs exhibited high anti-tumor efficacy, and significantly enhanced long-term *in vivo* tumor clearance in leukemia-bearing NSG mice as compared with conventional CD19-28ζ CAR-T. Mechanistically, CAPs were activated in an Lck-independent manner and displayed slower phosphorylation kinetics and a longer duration of signaling compared with 28ζ-CAR. The unique signaling properties of CAPs may therefore be harnessed to improve the *in vivo* efficacy of T cells engineered to express an anti-tumor chimeric receptor.

## Introduction

Chimeric antigen receptors (CARs) are molecules composed of an antibody fragment specific for a tumor antigen, fused to a transmembrane domain, a costimulatory domain, and a T-cell-signaling moiety, typically the T cell receptor zeta (TCRζ) chain. Most notably, CARs have been transformative in eradicating lymphoma, leukemia, and multiple myeloma (*1–4*). However, CAR efficacy in treating solid tumors has been limited. CAR-T cell exhaustion, cytokine-mediated toxicity, and disease relapse in situations where there is a low density of target antigen are several challenges for the successful use of CAR-T immunotherapy (*5*). As such, improvement in current CAR designs is a critical parameter that may be targeted to increase their efficacy.

CARs have been designed based on an attempt to harness TCR signaling. However, despite many permutations, CARs remain significantly less sensitive than TCRs (*6*). Unlike TCRs that can trigger T cell activation after the binding of as few as 1 to 10 ligands––a complex of agonist peptide and a molecule encoded by the major histocompatibility complex (pMHCs) (*7–9*), CARs require thousands of surface antigen molecules for productive signaling. Moreover, recent studies comparing TCR and CAR signaling revealed a blunting of proximal signaling from CARs (*6, 10, 11*). In an attempt to overcome these issues, we designed chimeric molecules with altered intracellular signaling domains. We hypothesized that incorporation of downstream T cell signaling molecules into the CAR design will allow for more sensitive and robust signaling. The proposed recombinant Chimeric Adapter Protein (CAP) is designed to bypass the TCRζ chain used in current FDA-approved CARs. Our original CAPs are chimeric molecules in which the extracellular targeting domain is linked to adapter domains in tandem with the kinase domain of ZAP70, a critical T cell protein tyrosine kinase (PTK).

The generation of CAPs was based on observations that following T cell activation, downstream adapter molecules form a distinct signaling cluster that segregates from the TCR complex and the kinase ZAP-70 (*12*). From this observation, we hypothesized that first, by linking the adapter proteins to ZAP70, an active adapter cluster would be generated by bypassing the need for TCR subunit activation. Second, direct triggering of the downstream signaling cascade via CAPs would bypass early events at the TCR, which behave as proofreading steps. These events are postulated to be required for crossing signaling thresholds before physiological T cell activation can be achieved (*13*). Thus, this type of bypass could potentially lead to a more sensitive and potent activation of T cells.

Here, we have designed and generated novel CAPs that bypass the TCRζ domains used in current FDA-approved CAR designs. These CAPs fuse an extracellular targeting domain to intracellular domains derived from downstream T cell signaling proteins that we have identified in distinct signaling clusters. CAPs harboring an scFv against CD19 (FMC63) and fused to LAT or SLP76 adapter moieties in tandem with the ZAP70 kinase domain, were generated. Importantly though, T cells expressing CAPs (CAP-Ts) containing adapter moieties promoted high levels of basal cytokine secretion in an antigen-independent manner. Therefore, CAPs that exclusively contained ZAP70 domains were further developed and these constructs demonstrated low basal activation and high antigen-specific cytokine production and cytotoxicity. First generation CAPs containing ZAP70 domains, and second-generation CAPs, containing ZAP70 and CD28 costimulatory domains, were further evaluated for their ability to eliminate CD19^+^ leukemia in a humanized NOD/scid/gamma (NSG) murine xenograft model. Second generation CAPs exhibited high anti-tumor efficacy, and significantly enhanced long term *in vivo* tumor clearance in leukemia-bearing NSG mice as compared with conventional CD19-28ζ CAR-T.

The enhanced efficacy of CAPs was associated with distinct signaling properties. Confocal and TIRF microscopy revealed a delayed recruitment of signaling molecules to CAP microclusters together with a decreased magnitude of signaling at the CAP immune synapse. Importantly, CAP signaling was maintained for a longer duration than 28ζ -CAR signaling. Moreover, the kinetics of activation of proximal signaling molecules, as evaluated as by determining their phosphorylation status was prolonged with CAP-induced signaling, We also found that CAPs but not CARs were activated in an Lck-independent manner. Thus, the increased tumor clearance and persistence by CAP-Ts may be due to the direct downstream activation of signaling molecules, bypassing inhibitory signals and resulting in a lower level but longer duration of signaling.

## Results

### CAPs can bypass upstream proteins and signal to downstream proteins

CAPs are chimeric molecules that contain adapter domains in tandem with the kinase domain of ZAP70, fused to an extracellular targeting domain. CAPs are designed to enable adapter phosphorylation by ZAP70 upon engagement of the extracellular domain. Adapter phosphorylation should then lead to activation of downstream signaling cascades and T cell activation, bypassing the TCRζ chain used xin current FDA-approved CARs **(Fig. 1A)**.

**Figure 1.**
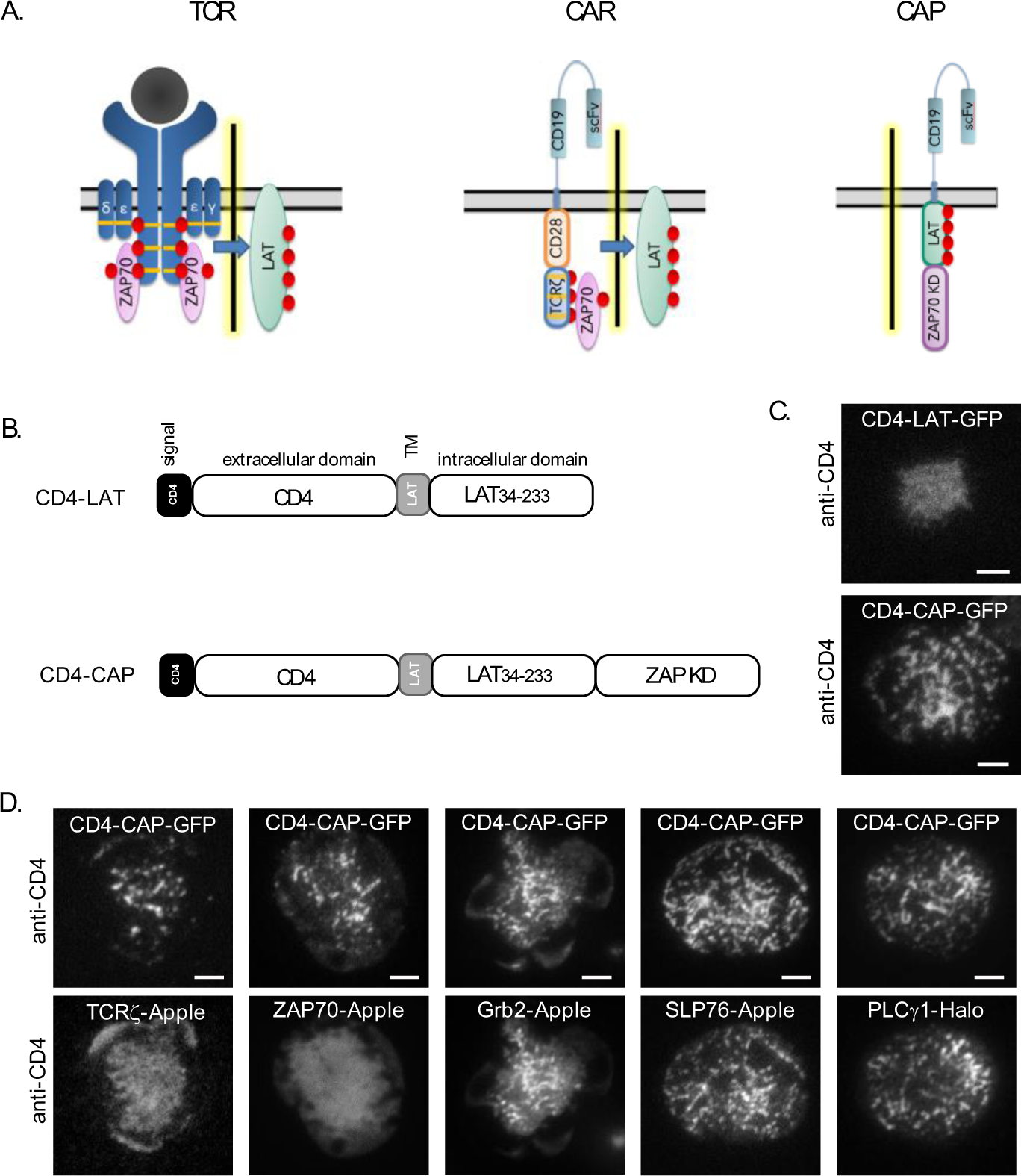
Chimeric Adapter Proteins (CAPs) can bypass upstream proteins and signal to downstream proteins. **A**. Schematic of TCR signaling, CAR signaling, and CAP signaling. While TCR and CAR must cross the signaling threshold (indicated as yellow highlighted bar) for productive T cell activation, CAP bypasses upstream steps. **B.** Schematics of CD4-LAT and CD4-CAP constructs. **C.** TIRF images of microclusters formed in Jurkat T cells expressing CD4-LAT-GFP or CD4-CAP-GFP activated on coverslips coated with anti-CD4 antibody. **D.** TIRF images of microclusters formed in Jurkat T cells expressing CD4-CAP-GFP (top row) and indicated fluorescent proteins (bottom row) activated on coverslips coated with anti-CD4 antibody. Scale bars in images, 2 μm.

We first tested CAP potential using the adapter protein LAT. As a proof-of-principle, a chimeric CD4-LAT construct was used as a backbone (*14*), fusing the C terminus to the ZAP70 kinase domain (KD) to generate CD4-CAP **(Fig. 1B)**. GFP tags were incorporated at the C terminus of both constructs and GFP-tagged CD4-LAT and CD4-CAP were expressed in Jurkat T cells. Consistent with previously published results (*14*), we observed that CD4-LAT-GFP did not cluster on anti-CD4 coated coverslips. In contrast, when CD4-CAP-GFP-expressing cells were dropped on anti-CD4 coated coverslips, CD4-CAP displayed robust microcluster formation and cell spreading, indicative of activation **(Fig. 1 C)**. These microclusters colocalized with phosphotyrosine **(fig. S1A)**, confirming an initiation of T cell activation. Microclusters did not form on anti-CD43 or anti-CD45 coated coverslips, indicating that binding of the CD4 extracellular domain to anti-CD4 antibodies specifically mediated cluster formation **(fig. S1B)**. CD4-CAP-GFP microclusters did not colocalize with TCRζ and ZAP70, but colocalized with downstream signaling proteins including Grb2, SLP76 and PLCγ1 **(Fig. 1D)**, cytosolic proteins that are recruited to phosphorylated LAT molecules upon TCR-mediated activation (*15*). These results indicate that in an *in vitro* Jurkat model, engineered CAP molecules bypass the TCRζ chain and signal to downstream proteins in a ligand-specific manner.

### Screening of CD19-CAP constructs containing LAT, SLP76 and ZAP70 domains

We next generated CD19-CAPs by replacing the CD4 extracellular domain of CD4-CAP with the anti-CD19 FMC63 moiety to generate CD19-CAP1 **(Fig. 2A)**. The LAT hinge and TM domains were also replaced with CD28 hinge and TM domains used in the FDA-approved second-generation CD19-28ζ CAR (*16*). To evaluate localization of CD19-CAP1 and assess whether CD19 binding led to intracellular signals, CD19-CAP1-GFP was generated and co-transfected with Grb2-Apple into Jurkat T cells. Upon interaction with CD19-expressing Raji B cells, both CD19-CAP1-GFP and Grb2-Apple showed robust recruitment to the immune synapse **(Fig 2B)**, indicating that CD19-CAP1 can signal to downstream proteins in an antigen-dependent manner. However, Jurkat T cells transfected with CD19-CAP1 showed high levels of basal CD69 expression compared with mock transfected controls **(fig. S2A)**, indicating that expression of CAP1 molecules cause high levels of tonic signaling. Though tonic signaling in T cells via the endogenous TCR and self-peptide loaded MHCs is associated with homeostasis (*17*), tonic signaling in CAR-Ts has been associated with adverse effects such as CAR-T exhaustion, limiting efficacy (*18*). In an attempt to decrease the tonic signaling observed in CAP1, we designed CAPs in which the ZAP70 Interdomain B (IB) domain was included, because fusion of the ZAP70 IB domain with the kinase domain (KD) has been shown to regulate ZAP70 kinase domain activity (*19*). CD19-CAP2 included the ZAP70 IB+KD fused with LAT. In CD19-CAP3, the LAT intracellular domain was replaced with SLP76, based on previous observations that the SLP76 adapter is capable of fully reconstituting LAT-deficient Jurkat T cells (*20*). In CD19-CAP3, the CD28 hinge and transmembrane domains were replaced with LAT sequences. Finally, CD19-CAP4, including only the ZAP70 IB+KD domains, was generated **(Fig. 2A)**. Cell surface expression of these constructs was assessed in primary human T cells and a construct similar to the FDA-approved CD19-4-1BBζ CAR was used as a positive control. While surface expression of all CAP constructs was significantly lower than that of the 4-1BBζ CAR positive control, differences in the relative cell surface expression of the different CAP constructs were also detected. CAPs that contained either LAT intracellular domains in tandem with ZAP70 domains (CAP1 and CAP2) or LAT TM domain and SLP76 intracellular domain in tandem with ZAP70 domains (CAP3) showed poor cell surface expression. However, CAP4, containing only intracellular ZAP70 domains, showed the highest surface expression amongst the tested CAP constructs **(fig. S2B and C)**.

**Figure 2.**
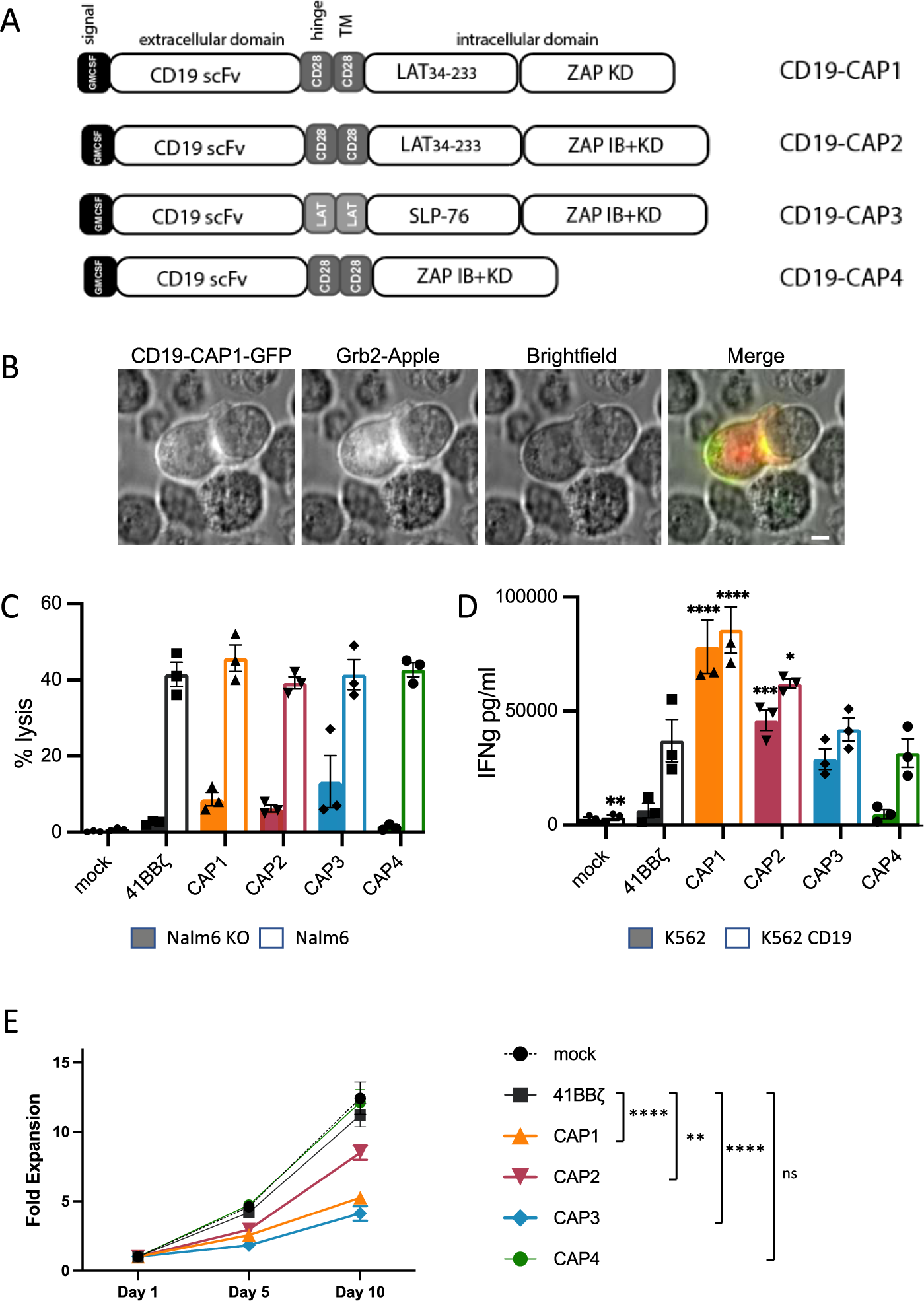
Screening of CD19-CAP constructs containing LAT, SLP-76 and ZAP-70 domains. **A**. Schematic of CD19-CAP constructs. **B.** Image of a Jurkat T cell expressing indicated fluorescent proteins forming an immune synapse with a Raji B cell. Scale bar in image, 2 μm. **C.** % lysis of target cells by control and CD19-CAP-expressing T cells incubated with indicated target cells at a 5:1 ratio. **D.** IFNγ production evaluated by ELISA from supernatants of control and CD19-CAP-expressing T cells incubated with the indicated target cells at a 1:1 ratio for 16 hrs. **E.** Proliferation of control and CD19-CAP expressing T cells. Two-way Anova analysis was performed in D and E comparing CAPs with 4-1BBζ control. Bars denote ±SEM. ns: P > 0.05. **: P ≤ 0.01; ****: P ≤ 0.0001. Data are representative of 3 independent experiments.

In an attempt to elucidate the characteristics that may have resulted in changes in expression levels and function of the different CAP constructs, we performed structural modeling. We found that the modification of the intracellular motif had a measurable influence on the properties of the CAP models, mainly affecting transmembrane (TM) domain stability. The major difference predicted by the modeling was that any molecules containing LAT sequences (CAP1, CAP2, CAP3) were unstable and failed to reach equilibrium using Molecular Dynamics. CAP4 was more stable (for details on protocols of model generation go to Supplemental Methods).

The functionality of CAPs were then compared with a 4-1BBζ-CAR in standard *in vitro* assays, evaluating cytotoxic activity and cytokine production. In a standard 4-hour cytotoxicity assay, CARs and CAPs showed nearly equivalent tumor cell killing **(Fig. 2C)**. Of note, cells expressing CAP1, CAP2 and CAP3 displayed slightly elevated antigen-independent cytoxicity, albeit <15%. In a standard overnight cytokine assay, T cells expressing CAP1, CAP2 and CAP3 produced significantly higher levels of IFNγ than the 4-1BBζ positive control CART cells. Notably, antigen-independent IFNγ production was also high in these groups indicating a high level of tonic signaling by these CAPs. CAP4 was the only CAP molecule amongst those tested that showed robust cytokine production, comparable with 4-1BBζ CAR T cells in a strictly antigen-dependent manner **(Fig. 2D)**. Moreover, cells expressing CAP4 displayed robust proliferation, comparable with mock-transduced and 4-1BBζ CAR T cells **(Fig. 2E)**. This screening of CAP constructs indicates that CAP4 is the only CAP that is highly functional in an antigen-dependent manner. Therefore, we focused our further development of CAP constructs on the CAP4 backbone, which contained only ZAP70 domains in the intracellular region.

### Screening of CD19-CAP constructs containing ZAP70 domains

CAP4-modified constructs all contained the CD19scFv extracellular domain, the CD28 hinge and TM domain, and various ZAP70-containing intracellular domains **(Fig. 3A)**. The CAP4.2 construct is identical to the original CAP4 construct but contains a G_4_S linker between the CD28 TM domain and ZAP70-IB+KD domain, potentially affording more flexibility for the intracellular domain. Additionally, we designed CAP4 constructs that contained the CD28 costimulatory domain. In CAP4.6, the CD28 costimulatory domain was fused with the ZAP70 IB+KD domain. We also generated CAP4.7, which includes full-length ZAP70, including the N-terminal SH2 domains, because the N-terminus of ZAP70 plays an important role in regulating the threshold of T cell signaling (*21–23*). Finally, CAP4.8 was designed to include a mutated CD28 signaling domain that cannot bind downstream signaling proteins (*24*). The four CAP4 constructs thus fall into two categories: generation 1 CAPs (CAP4.2 and CAP4.8) that do not contain costimulatory capacity, and generation 2 CAPs (CAP4.6 and CAP4.7) that have functional CD28 costimulatory capacity. Expression of these constructs was tested in T cells with CD19-4-1BBζ CAR as a positive control. Surface expression of all CAP4 constructs, except for CAP4.7, which includes full-length ZAP70 and is significantly larger than all other constructs tested, were similar. While CAP4.2, CAP4.6 and CAP4.7 expression were 25-30% lower than the 4-1BBζ-CAR, CAP4.7 expression was ∼70% lower than the positive control **(fig. S3A and B)**. Evaluation of total cellular expression of these constructs by western blotting of whole cell lysates under reducing conditions showed expected mobilities **(fig. S3C-E)**. Moreover, when expression of the constructs in whole cell lysates was detected under non-reducing conditions without DTT to evaluate oligomerization, we detected higher order protein complexes of CARs and CAPs (**fig. S3C-E).** These data suggest that these chimeric molecules form covalently-linked oligomers in cells under non-stimulated conditions.

**Figure 3.**
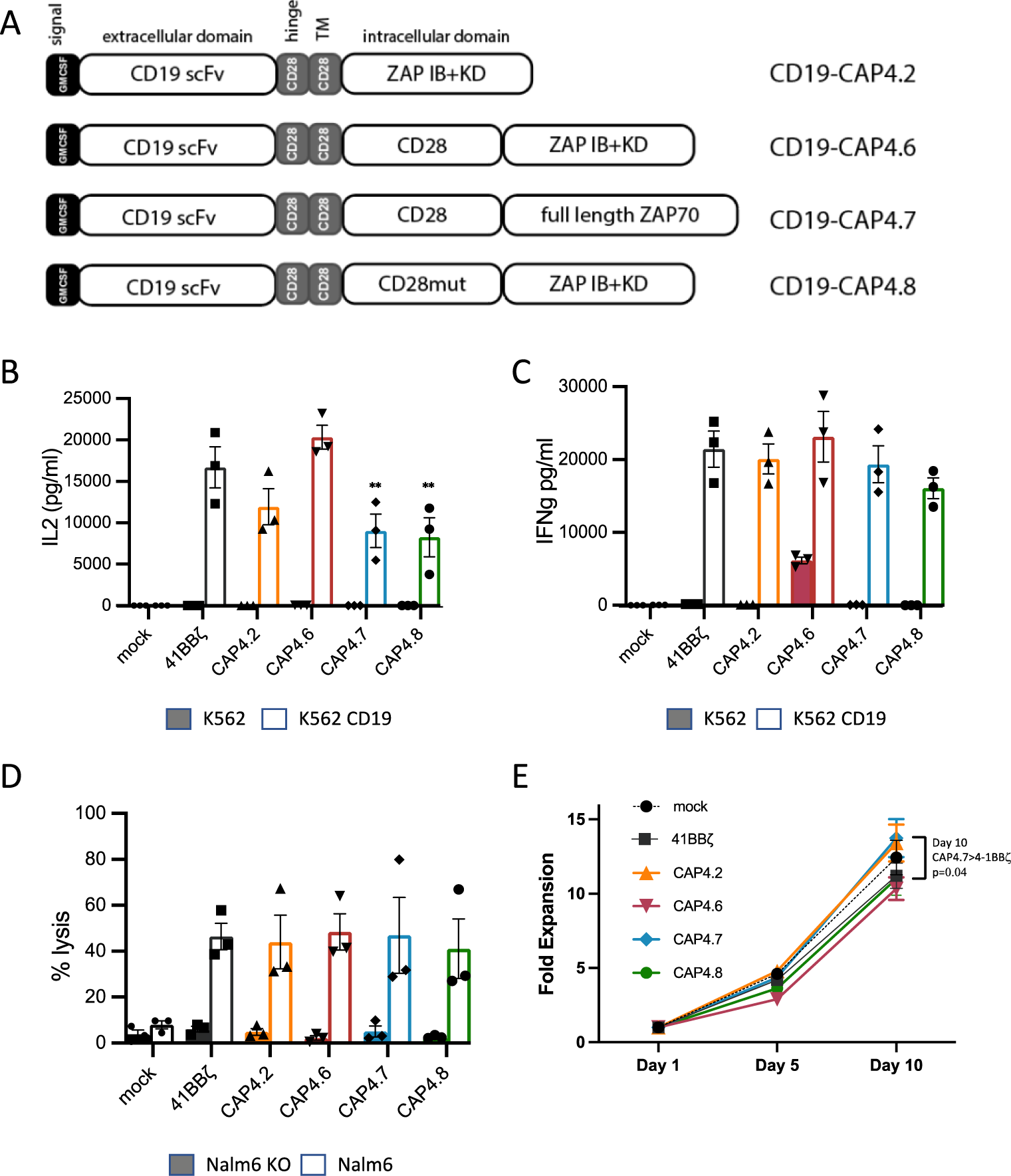
Screening of CD19-CAP4 constructs that contain ZAP-70 domains. **A**. Schematic of CD19-CAP4 constructs. **B and C.** IL2 and IFNγ production by control and CD19-CAP4 expressing T cells incubated with indicated target cells at a 1:1 ratio for 16 hrs. **D.** % lysis of target cells by control and CD19-CAP4 expressing T cells incubated with indicated target cells at a 5:1 ratio. **E.** Proliferation of control and CD19-CAP4-expressing T cells *in vitro* as a function of time. Two-way Anova analysis was performed comparing CAPs with 41BBz control. Bars denote ±SEM. *: P ≤ 0.05. Data are representative of independent experiments from 3 different donors.

The functionality of CAPs were then compared with a 4-1BBζ-CAR in standard *in vitro* assays, evaluating cytotoxic activity and cytokine production. In a standard overnight cytokine assay, all CAPs produced IL2 at levels comparable to the 4-1BBζ positive control CAR in response to target antigen **(Fig. 3B)**. While CAP4.6-expressing T cells produced elevated basal levels of IFNγ, all other CAPs produced IFNγ at levels comparable with the 4-1BBζ positive control CAR in a target antigen-dependent manner **(Fig. 3C)**. In a standard 4-hour cytotoxicity assay, CARs and all CAP4s showed equivalent tumor cell killing in a strictly antigen-dependent manner **(Fig. 3D)**. Finally, T cells expressing CAP4s displayed robust proliferation, comparable with mock-transduced and 4-1BBζ CAR-expressing T cells **(Fig. 3E)**. Together, our *in vitro* analyses indicate that addition of the CD28 costimulatory domain only increased antigen-independent cytokine secretion in the presence of the ZAP70 IB+KD domains alone. All other CAP4 constructs were highly functional in an antigen-dependent manner.

### CD19-CAP4 constructs show robust efficacy in an *in vivo* NSG leukemia model

In order to further differentiate CAP4 candidates, we evaluated their *in vivo* efficacy in an immunodeficient NOD/scid/gamma (NSG) mouse model **(Fig. 4A)**. As high levels of signaling by 28ζ-CARs are linked to T cell exhaustion in the setting of high antigen density (*11, 25, 26*), we tested sub-curative doses of CAP4 versus 28ζ -CAR T cells (3e6) in NSG mice engrafted with Nalm6 leukemia cells that express high levels of CD19. Surface expression of 28ζ-CAR and CAPs were evaluated on the transduced donor T cells prior to infusion in NSG mice. CAPs 4.2, 4.6 and 4.8 showed similar levels of expression to 28ζ-CAR, but surface expression of CAP4.7 was ∼45% lower than the positive control (**fig. S4A).** While the percentages of naïve, central memory and effector T cell subsets were similar in all groups, 28ζ-CAR transduced donor T cells exhibited higher levels of the CD25 (IL2Rα) activation marker as compared to T cells transduced with the different CAP constructs **(fig. S4B and C)**. These data suggest that CAP-transduced T cells may have a lower level of basal signaling than 28ζ-CAR T cells.

**Figure 4.**
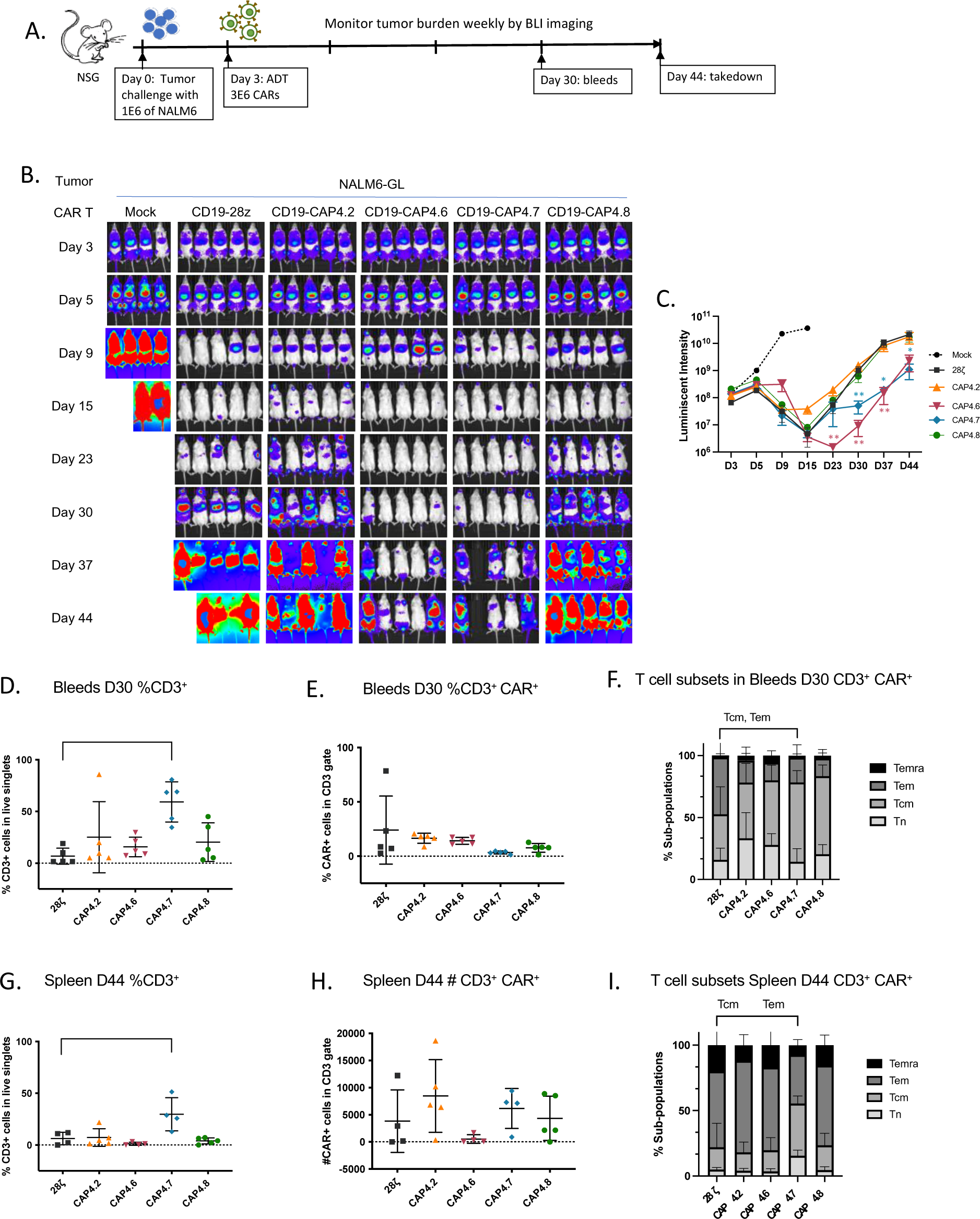
CD19-CAP4 constructs show robust efficacy in an *in vivo* NSG leukemia model. **A**. Schematic of NSG mouse model of leukemia. Luciferase-transduced NALM6 cells (1×10^6^) were injected intravenously via tail vein into NSG mice on day 0. Engraftment was documented by bioluminescent imaging (BLI) on day 3 and cohorts of five mice were randomized to intravenous treatment with mock transduced T cells, 28-ζ CAR-Ts or one of the CD19-CAPs as designated (3×10^6^ CAR^+^ or CAP^+^ cells/mouse). Mice were followed by weekly BLI. **B.** Leukemia growth was evaluated at the indicated timepoints by bioluminescent imaging (BLI) and IVIS images images are shown. **C.** Quantification of the BLI radiance data for each individual mouse is presented. Bars denote ±SEM. Statistical differences were assessed using a Mann Whitney t-test comparing CAP4.6 or CAP4.7 with 28z-CAR. *: P ≤ 0.05; **: P ≤ 0.01. **D-F.** Flow cytometric analysis of peripheral blood on Day 30 showing the percentages of human CD3^+^ T cells (**D**), CAR^+^ T cells (**E**), and T cell subsets (**F**). **G-I.** Flow cytometric analysis of splenocytes on Day 44 assessing CD3^+^ T cells (**G**), CAR+ T cells (**H**), and T cell subsets as follows: T_n_ (CD62L^+^CD45RA^+^), T_cm_ (CD62L^+^, CD45RA^-^), T_em_ (CD62L^-^CD45RA^-^) and T_emra_ (CD62L^-^CD45RA^-^) (**I**). **D-I.** Statistical differences were assessed using a two way Anova. Bars denote ±SEM. ns: P > 0.05; *: P ≤ 0.05; **: P ≤ 0.01

28ζ -CAR-Ts and all tested CAP4-Ts exhibited early efficacy in reducing tumor burden as compared with mock-transduced T cells **(Fig. 4B and C)**. Notably though, by day 30, mice treated with 28ζ -CAR, CAP4.2, and CAP4.8 relapsed, while CAP4.6 and CAP4.7-treated mice achieved a more durable tumor regression. Flow cytometry analyses of peripheral blood T cells at 30 days following tumor injection revealed a higher percentage of CD3^+^ T cells in CAP4.7-treated mice as compared with 28ζ -CAR positive control **(Fig. 4D)**. Interestingly, detectable surface CAR and CAP expression in all groups was low (means of <25%), potentially due to internalization after activation **(Fig. 4E)** (*27*). We also assessed the differentiation states of the adoptively transferred T cells, as phenotype has been shown to strongly correlate with antitumor potency (*28, 29*). Analyses of T cell subsets in peripheral blood revealed a higher percentage of central memory cells (T_cm_ CD62L^+^ CD45RA^-^), and fewer effector memory cells (T_em_ CD62L^-^ CD45RA^-^) in CAP4.7-Ts as compared with 28ζ - CAR-Ts **(Fig. 4F)**. These trends were also observed in flow analyses of spleen at Day 44 **(Fig. 4 G-I)**. Thus CAP4.7-Ts induce a more durable remission, associated with a more enhanced accumulation of central memory T cell populations compared with T cells transduced with the conventional 28ζ -CAR vector.

We next sought to compare CAP4 constructs with 4-1BBζ -CAR, because 4-1BBζ -CAR T-cells have been suggested to exhibit less exhaustion and better persistence than 28ζ -CAR T-cells (*18*). As seen in the previous experiment, surface expression of 4-1BBζ-CAR and CAPs 4.2, 4.6 and 4.8 were similar, with lower surface expression of CAP4.7 (**fig. S5A).** Donor T cells transduced with 4-1BBζ -CAR and CAPs showed similar levels of CD25 surface expression as well as similar percentages of naïve, central memory, and effector memory T cell subsets (**fig. S5B and C)**. 4-1BBζ-CAR and all CAPs tested were able to eradicated the engrafted leukemia, but tumors returned in mice treated with first generation CAP (CAP4.2 and CAP4.8). Notably though, we detected durable tumor control in mice treated with T cells transduced with 4-1BBζ -CAR and second-generation CAPs (CAP4.6 and CAP4.7; **fig. S5D and E**). Flow cytometry analyses of peripheral blood at Day 34, the time when tumor control began to diverge, showed that CD3^+^ cells were present at higher levels in 4-1BBζ -CAR and CAP4.7-treated mice compared to other CAPs (**fig. S5F)**. However, detectable surface CAP expression on all CAP groups was lower than the 4-1BBζ -CAR T-cell control (**fig. S5G)**. Analyses of T cell differentiation states in the spleen showed similar results and a higher percentage of T_cm_ and lower percentage of T_em_ in CAP4.7-Ts compared with 4-1BB ζ-CAR-Ts **(fig. S5H-J)**. Taken together, these *in vivo* experiments demonstrate that second generation CAPs exhibit efficacy, mediating durable remissions of Nalm6 leukemia. Moreover, CAP4.7-Ts despite having the lowest expression of a CAP molecule at the initiation of the *in vivo* experiment, have the best expansion and least differentiated profile compared with both 28ζ and 4-1BBζ -CAR T cells.

We conducted structural modeling to gain insights into the functional differences between the CAP4 constructs. Molecular models were generated using a previously described protocol (*30*). The results of this analysis suggest that minor changes in the intracellular domain significantly impact the transmembrane domain. CAP4.2 appears to be less stable due to the GGGS linker while the inclusion of the CD28 intracellular domain appears to have a stabilizing effect in CAP4.6, CAP4.7 and CAP4.8. While a quantitative comparison of the models is difficult due to the lack of experimental structural information for full-length CAR-Ts, we explored the use of AlphaFold as a means of generating partial models of the different CAPs. AlphaFold proved helpful for the generation of ZAP70-based dimers. We used these statistically better-predicted regions and combined them with our previous models to generate suitable chimeras to further assess the effect of sequence modifications on CAR stability. Renditions of the models are presented in **fig. S6**. The Buried Solvent Accessible Area value (BSAS), which correlates with molecule stability, varies widely for the CAP4 series, with the CAP4.2 construct exhibiting the lowest BSAS value of 549.8Å^2^. CAP4.6, CAP4.7, and CAP4.8 all had higher BSAS values, with CAP4.7 having the highest BSAS value (1372.3 Å^2^, **fig. S6F**). Importantly, BSAS values correlated with the experimental data, with CAP4.7 appearing to exhibit the most stability, comparable to previously reported data for FMC-63 28ζ and Hu19-CD8-28ζ CARs (*30*).

### Immune Synapses and microclusters generated by CAP4.7 and 28-ζ constructs show distinct properties

To begin to assess the mechanisms accounting for the efficacy of second generation CAPs (CAP4.6 and CAP4.7), we first evaluated their subcellular localization by microscopy in the context of the immunological synapse (IS) and microclusters. Lentiviral constructs encoding GFP-tagged 28ζ -CAR and CAP4.7 constructs were generated and expressed in Jurkat T cells harboring ZAP70-Apple. These Jurkats were incubated on coverslips loaded with CD19-expressing Raji B cells. Cell conjugates were then fixed after 10 minutes and immunostained for phospho-SLP76 (pSLP76) to evaluate proximal signaling. This method allowed us to visualize both the chimeric molecules and proximal signaling proteins at the IS. Robust recruitment of both 28ζ - CAR and CAP4.7 at the IS was observed at this time point, with equivalent synapse volumes of CAR and CAP **(Fig. 5A and B)**. By contrast, ZAP70 and pSLP76 were present at significantly lower levels in the CAP4.7 synapse **(Fig. 5C)**. When intensity at the IS was normalized to the total cellular intensity of the corresponding protein, pSLP was still recruited at significantly lower levels at the CAP4.7 IS **(Fig. 5D)**. Thus, CAP-expressing cells efficiently form synapses and recruit CAP molecules to the IS at similar levels to 28ζ-CAR, albeit with recruitment of lower levels of phosphorylated proximal signaling molecules at CAP synapses.

**Figure 5.**
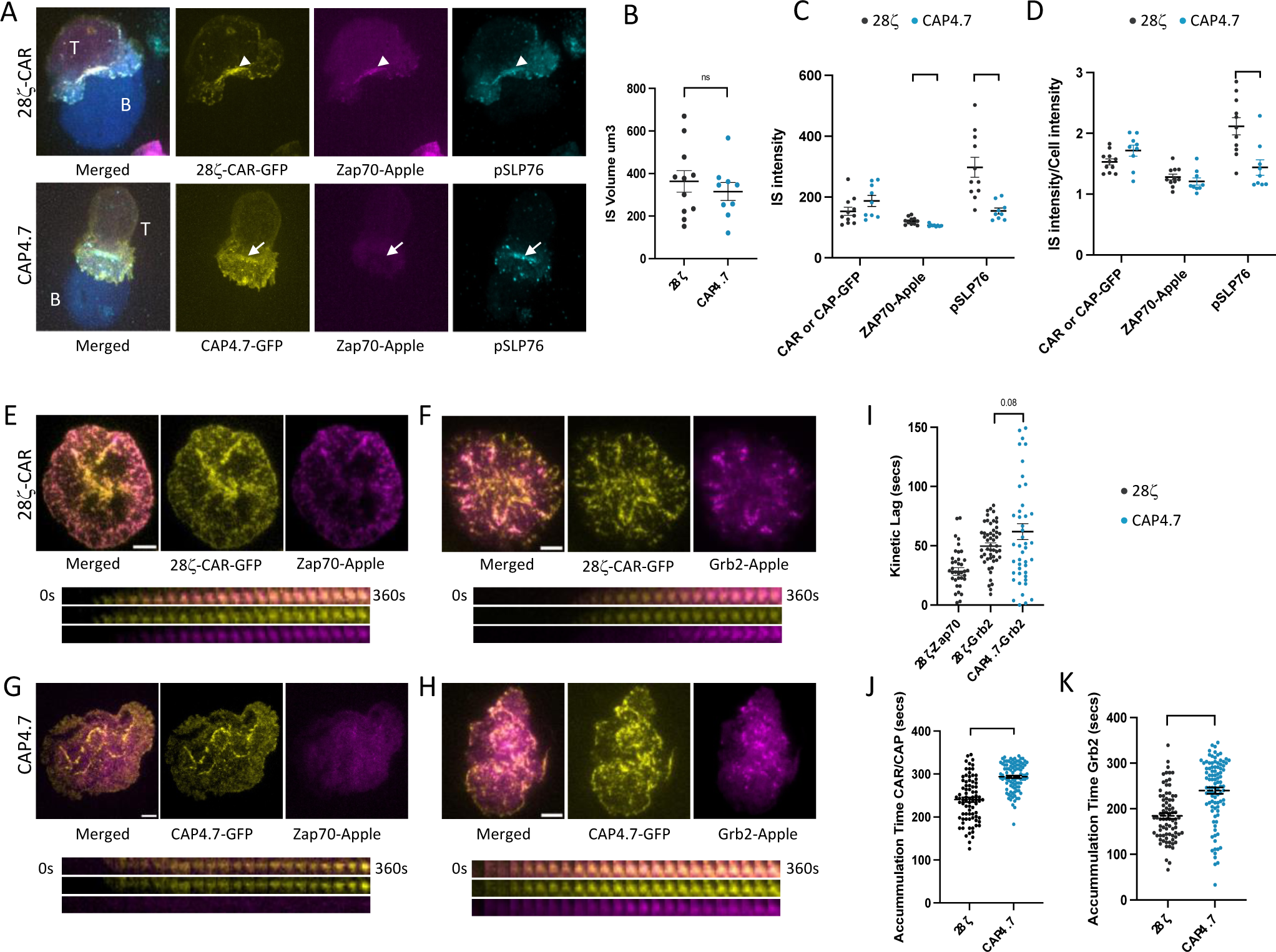
CAP-T cells display lower recruitment of activated proteins and delayed and prolonged kinetics of protein recruitment than CAR-T cells. **A.** Fixed cell images of a 28ζ-CAR-GFP cell (above; GFP is pseudo colored in yellow) or CAP-GFP expressing cells (below) interacting with a Raji B-cell (blue). CAR and CAP expressing cells were also transfected with ZAP70-Apple (magenta) and immunostained for pSLP76 (turquoise). **B.** Volume of the IS formed by 28ζ-CAR or CAP4.7 expressing cells. **C.** A comparison of fluorescence intensity at the immune synapse for the indicated proteins. **D and E.** TIRF images of 28ζ-CAR-GFP (yellow) and Zap70-Apple (magenta; D) or Grb2-Apple (magenta; E). Below is a time-lapse montage showing a single microcluster at 15 s/frame over the course of 6 minutes. **F and G.** TIRF images of CAP4.7-GFP (yellow) and Zap70-Apple (magenta; F) or Grb2-Apple (magenta; G), below is a time-lapse montage showing a single microcluster at 15 s/frame over the course of 6 minutes. **H.** The kinetic lag times that are calculated from the half-max intensity of the best fit sigmoidal curve of the fluorescence intensity of each microcluster. **I and J.** Accumulation time, which is how long it takes for the CAR or CAP (I) or Grb2 (J) to accumulate until maximum fluorescence intensity is reached. Welch’s t-test was performed. Dot plots show mean ± SEM. ns: P > 0.05; **: P ≤ 0.01; ****: P ≤ 0.0001. Data are representative of 3 independent experiments.

To investigate whether the lower amount of proximal signaling proteins at CAP synapses was due to a change in recruitment kinetics, we next evaluated recruitment of signaling molecules in live cell imaging experiments at CAR/CAP microclusters. We transfected GFP-tagged CAR and CAP cells with ZAP-Apple or Grb2-Apple and evaluated recruitment of Apple-tagged proteins to GFP microclusters formed on CD19-Fc-coated coverslips by TIRF microscopy and imaged cells at room temp (21 °C), to slow down microcluster formation kinetics. Quantification of fluorescent intensities showed that ZAP and Grb2 were recruited to 28ζ-CAR microclusters with similar kinetics (∼30sec for ZAP70 to 28ζ-CAR and ∼60 sec for Grb2 to 28ζ-CAR) as previously reported for recruitment to TCRζ (*12*) **(Fig. 5E and F)**. ZAP-Apple recruitment to CAP microclusters was not observed, consistent with reduced recruitment of ZAP70 to the CAP IS **(Fig. 5G)**. Two additional differences were observed at CAP microclusters. First, recruitment of Grb2 molecules to CAP microclusters was delayed as assessed from the kinetic lag measurements of the differences in half-max fluorescent intensities **(Fig. 5H and I)**. The difference between CAR and CAP in recruitment kinetics of Grb2 becomes even more apparent when the CAP-Grb2 kinetic lag (61 sec) is compared with the corresponding ZAP-Grb2 kinetic lag in 28ζ−CAR expressing cells. This parameter can be extracted by subtracting the 28ζ-ZAP (28.26 sec) lag from the 28ζ-Grb2 lag (49.54) to yield a value of 21.28 sec for ZAP-Grb2, which is considerably shorter than the 61 sec lag observed for Grb2 to be recruited to the CAP. Second, CAP microclusters as well as Grb2 recruited to CAP microclusters, accumulated for significantly longer periods of time compared with 28ζ -CAR microclusters **(Fig. 5 J and K)**. Together these data indicate that while the magnitude of signaling at the CAP IS decreased and the kinetics of recruitment of signaling molecules to CAP microclusters is delayed, CAP signaling is maintained for a longer duration than 28ζ - CAR signaling.

### Signaling downstream of CAPs and 28ζ -CAR differ in strength and kinetics

To assess the extent to which CAPs engage the signaling machinery of the TCR complex relative to a CAR, the kinetics of activation of a panel of proximal signaling molecules were evaluated as a function of their phosphorylation status. Jurkat T cells that had been lentivirally transduced with 28ζ -CAR, CAP4.6 or CAP4.7 constructs were incubated with CD19-negative (parental) or CD19-positive K562 target cells. Phosphorylation of the chimeric receptors themselves was detected using a phospho-TCRζ antibody. 28ζ -CAR phosphorylation (pCAR) peaked early at 2 min and showed rapid dephosphorylation **(Fig. 6A and B)**. In contrast, CAP phosphorylation (pCAP) as detected by phospho-ZAP, showed slower and more stable phosphorylation kinetics. CAP4.6 had high basal rates of phosphorylation that increased upon stimulation, while CAP4.7 had no detectable basal signaling and absolute levels of phosphorylation were much lower than that of CAP4.6. Importantly though, phosphorylation kinetics were similar between both CAPs; they showed peak phosphorylation at 15 min after which both CAPs showed slow rates of dephosphorylation. Thus, rates of phosphorylation onset and dephosphorylation of CAPs were lower than for 28ζ-CAR **(Fig. 6C and D)** and CAPs remain phosphorylated for a longer duration.

**Figure 6.**
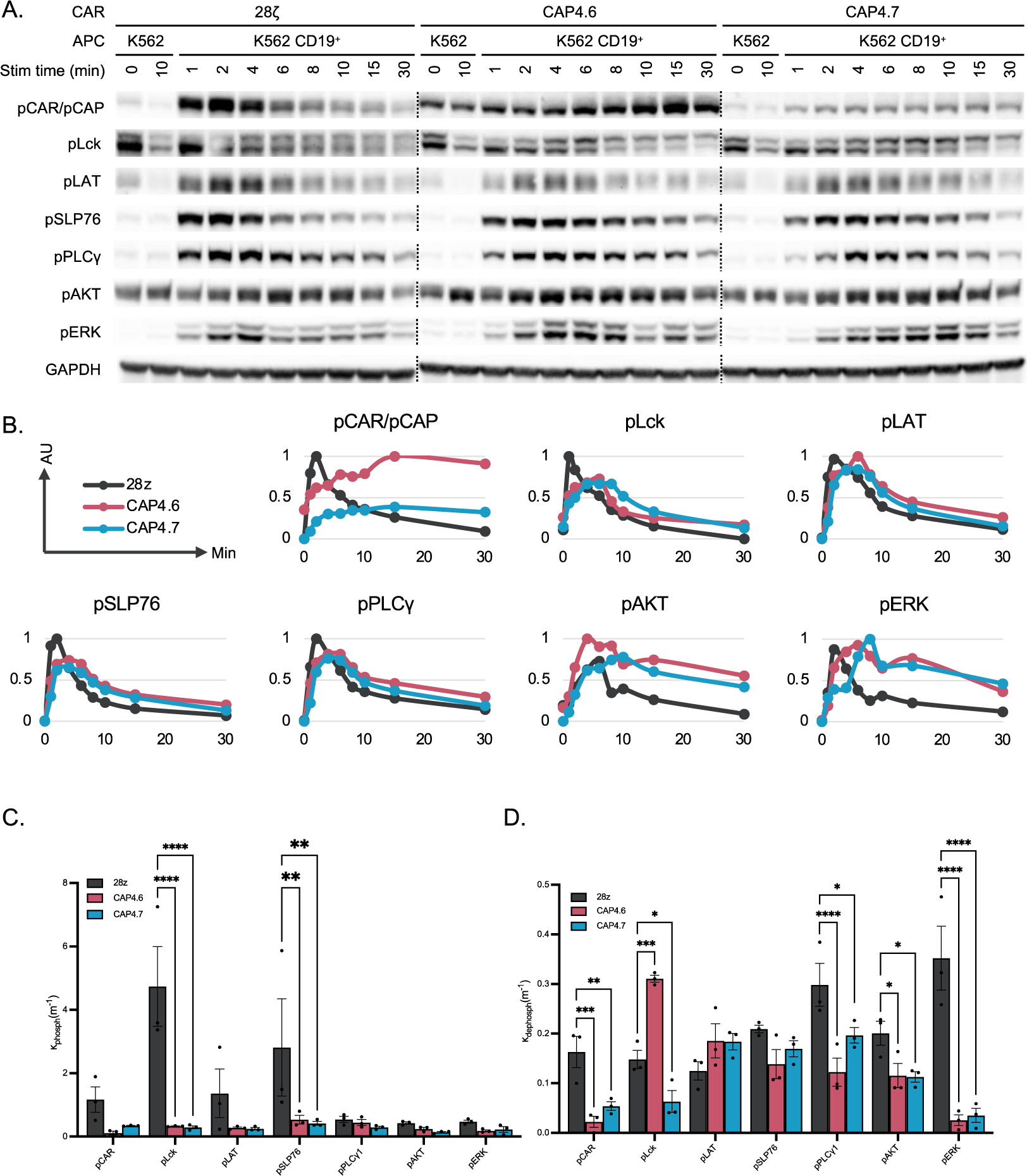
Phosphorylation Kinetics of CAR and CAPs. Jurkat-E6.1 cells stably expressing 28ζ-CAR or indicated CAP constructs were mixed with antigen presenting cells (APCs). APCs were either antigen negative (K562s) or antigen positive (K562s stably transduced with CD19). Cell mixtures were incubated at 37°C for given amounts of time and then lysed and immunoblotted for phosphorylated forms of immune signaling markers. Blot volumes were quantified using Bio-Rad Laboratories’ Image Lab software and normalized to total protein. V = Volume_pProtein_/Volume_GAPDH_. AU = (V-V_minimum_)/(V_maximum_-V_minimum_). **A.** Representative blots of 28ζ, CAP4.6, and CAP4.7 signaling kinetics. **B.** Averaged graphs of phosphorylation curves for different markers of CAR/CAP activation/signaling. The lower value of the two negative controls (K562 0min or 10min) was used for the 0min timepoint for each marker. **C and D.** Phosphorylation and dephosphorylation rates for signaling markers in each construct. Rates were determined by fitting an expression modeling exponential rise and decay. Data are representative of three independent experiments.

We next evaluated TCR proximal signaling events **(Fig. 6A)**. Both CAR and CAP molecules induced sequential phosphorylation of Lck, LAT, SLP76, PLCγ1, Akt and ERK, all of which are involved in the classic TCR signaling pathway. Similar to the phosophorylation data for the chimeric receptors themselves, signaling molecules remained phosphorylated for a longer duration in CAP-stimulated cells **(Fig. 6B)**. While signaling levels of the more proximal molecules (Lck, LAT, SLP76 and PLCγ1) were higher in 28ζ -CAR than CAPs, those of more distal molecules, Erk and Akt, were higher in CAPs and remained elevated for an extended time period **(Fig. 6B and fig. S7A)**. We also evaluated whether the signaling molecules included as components in the chimeric receptors (TCRζ for 28ζ -CAR and ZAP70 for CAPs) were phosphorylated in the cell. Unexpectedly, only CAP4.7 induced high levels of endogenous TCRζ phosphorylation **(fig. S7B and C)**. As expected, 28ζ-CAR induced high levels of endogenous ZAP70 phosphorylation, but surprisingly, a low level of endogenous ZAP70 phosphorylation was also observed downstream of CAPs **(fig. S7B and C)**. Together these data indicate that although CAPs show slower kinetics and lower magnitude of phosphorylation than 28ζ -CAR, the CAPs themselves as well as downstream signaling proteins remain activated for a longer duration. These signaling properties are consistent with the functionality of these cells: 28ζ -CAR exhibited strong effector function but decreased persistence, while second-generation CAPs showed better persistence and long-term tumor control.

### CAPs can propagate signals in the absence of Lck

To more fully elucidate the mechanism(s) of CAP activation, we next examined the requirements for proximal signaling molecules in the initiation and signaling of 28ζ-CAR and CAPs. To this end, we compared 28ζ-CAR and CAP signaling in Jurkat cell lines that were deficient in either Lck or ZAP70 expression. In the case of 28ζ-CAR, CAR phosphorylation itself was reduced by Lck deficiency, but not significantly affected by the lack of ZAP70. Interestingly though, phosphorylation of signaling molecules (ZAP70, LAT, SLP76, PLCγ1, Erk) downstream of 28ζ-CAR showed a higher dependence on ZAP70 than Lck, indicating the presence of Lck-independent CAR activation **(Fig. 7A and B)**. In comparison, in CAP-transduced cells, phosphorylation of the CAP molecules themselves (CAP4.6 and CAP4.7) as well as signaling of proximal signaling molecules did not require Lck. Surprisingly, loss of ZAP70 had a deleterious effect on phosphorylation of CAP4.7, as well as phosphorylation of downstream proximal molecules (LAT, SLP76, PLCγ1), indicating a requirement for endogenous ZAP70 in the activation of CAP4.7 **(Fig. 7A and B)**. While CAP4.6 also showed a partial requirement for endogenous ZAP70 in the phosphorylation of LAT, SLP76, and PLCγ1, signaling was maintained. Because a robust phosphorylation of endogenous TCRζ was detected downstream of CAP4.7 (**fig. S8A**), we investigated the requirement of endogenous TCR in CAP4.7 signaling by expressing 28ζ-CAR and CAP4.7 in a TCRβ KO Jurkat cell line. Neither 28ζ-CAR nor CAP4.7 showed a requirement for endogenous TCR **(Fig. 7A)**.

**Figure 7.**
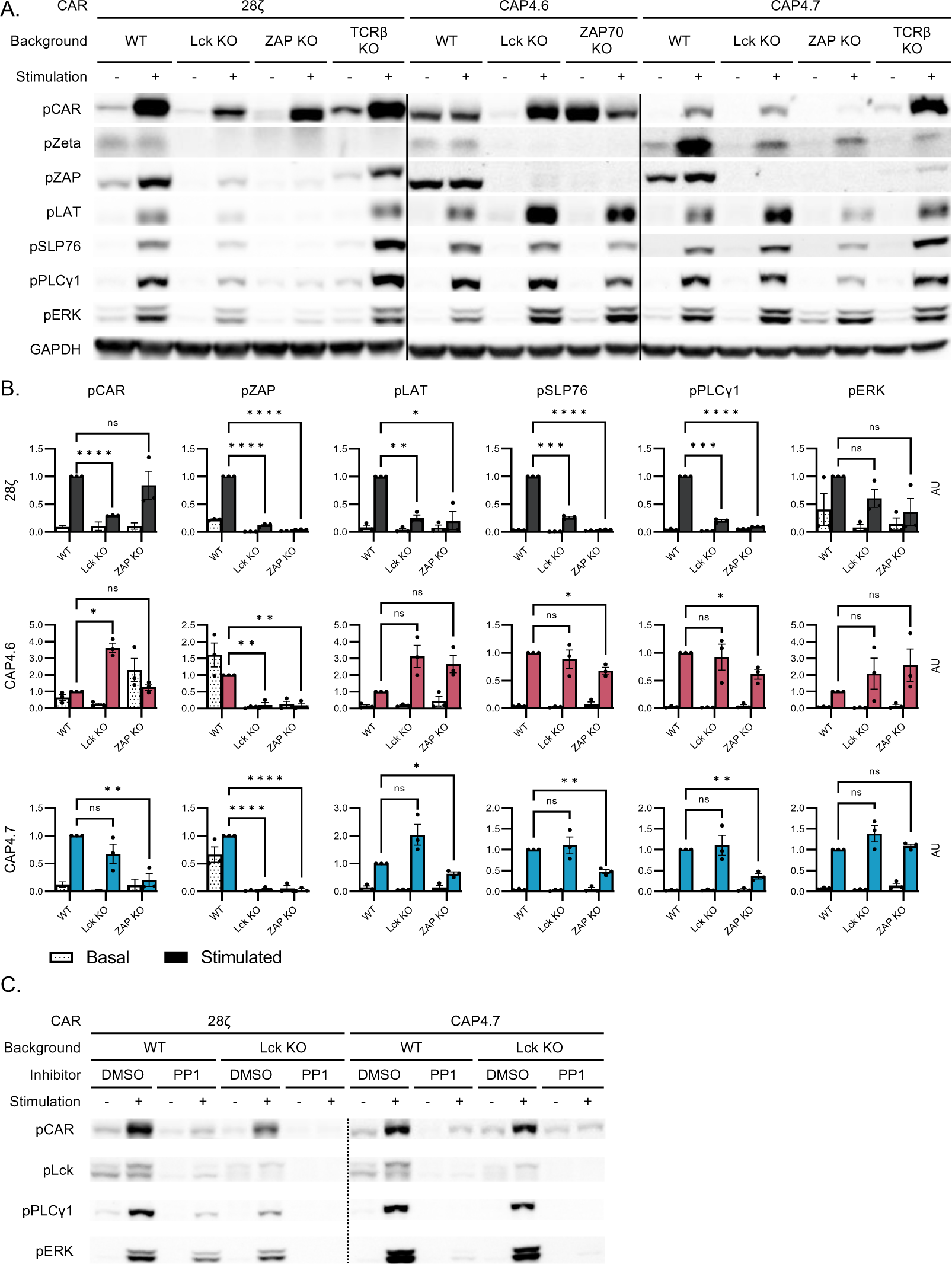
CAR and CAP Dependency on Proximal Immune Proteins. CAR or CAP constructs were stably transduced into Jurkat-E6.1 cells that either had Wild Type, CRISPR Lck KO, P116 ZAP KO, or CRSIPR TCRβ KO genetic background. Those cells were mixed with either antigen positive or antigen negative APCs, incubated at 37°C for 8 minutes, and then lysed and immunoblotted for phosphorylated forms of immune signaling markers. Blot volumes were quantified using Bio-Rad Laboratories’ Image Lab software and normalized to total protein. V = Volume_pProtein_/Volume_GAPDH_. AU = (V-V_minimum_)/(V_maximum_-V_minimum_). **A.** Representative blots of 28ζ, CAP4.6, and CAP4.7 signaling in the absence of either nothing, Lck, ZAP, or TCR. **B.** Averaged graphs of phosphorylation intensity for different markers of CAR/CAP activation/signaling. Paired Student’s T Tests were performed comparing the phosphorylation levels of activation/signaling markers in stimulated WT vs Lck KO or ZAP KO cells. Data are representative of three independent experiments. Bars denote ±SEM. ns: P > 0.05; *: P ≤ 0.05; **: P ≤ 0.01; ***: P ≤ 0.001. ****: P ≤ 0.0001. **C.** One blot of 28ζ and a representative blot of CAP4.7 signaling in the absence of either nothing or Lck and preincubation with either DMSO or PP1 (a Src family kinase inhibitor).

Low levels of 28ζ -CAR activation and normal levels of CAP activation in the absence of Lck was intriguing. To determine if any other Src family kinase (SFK) was responsible for initiation of Lck-independent 28ζ-CAR and CAP signaling, we used the pan-Src kinase inhibitor PP1. Upon PP1 treatment, 28ζ-CAR and CAP4.7 phosphorylation and downstream signaling were abrogated in both WT and Lck KO backgrounds **(Fig. 7C and fig. S8A)**. As Lck and Fyn are the main SFKs in T cells (*31*), these data strongly suggest that a Src family kinase, most likely Fyn, is responsible for Lck-independent 28ζ-CAR and CAP activation.

### Lck is a driver of CAR and CAP Degradation

In the course of our kinetics study of CAR and CAP phosphorylation in **Figure 6**, we observed that CAR and CAP expression were significantly downregulated following antigen recognition as has been previously reported for CAR-Ts (*27*). While studying CARs and CAPs in various mutant Jurkat cell lines, we also noted that degradation of 28ζ-CAR and CAPs did not occur to the same extent in Lck KO cells **(fig. S8B)**. It has been proposed that CAR-T cell persistence and functionality can be enhanced by blocking antigen-induced CAR degradation (*32*). Therefore, we examined 28ζ-CAR and CAP degradation upon antigen encounter in WT and Lck KO cells. Both 28ζ-CAR and CAP expression in WT Jurkat cells was decreased by 60% after encountering antigen in a 30 min time course. In contrast, degradation of 28ζ-CAR and CAP was significantly reduced in Lck-deficient Jurkat cells **(Fig. 8)**. These results point to Lck as a major driver of CAR and CAP degradation upon antigen encounter.

**Figure 8.**
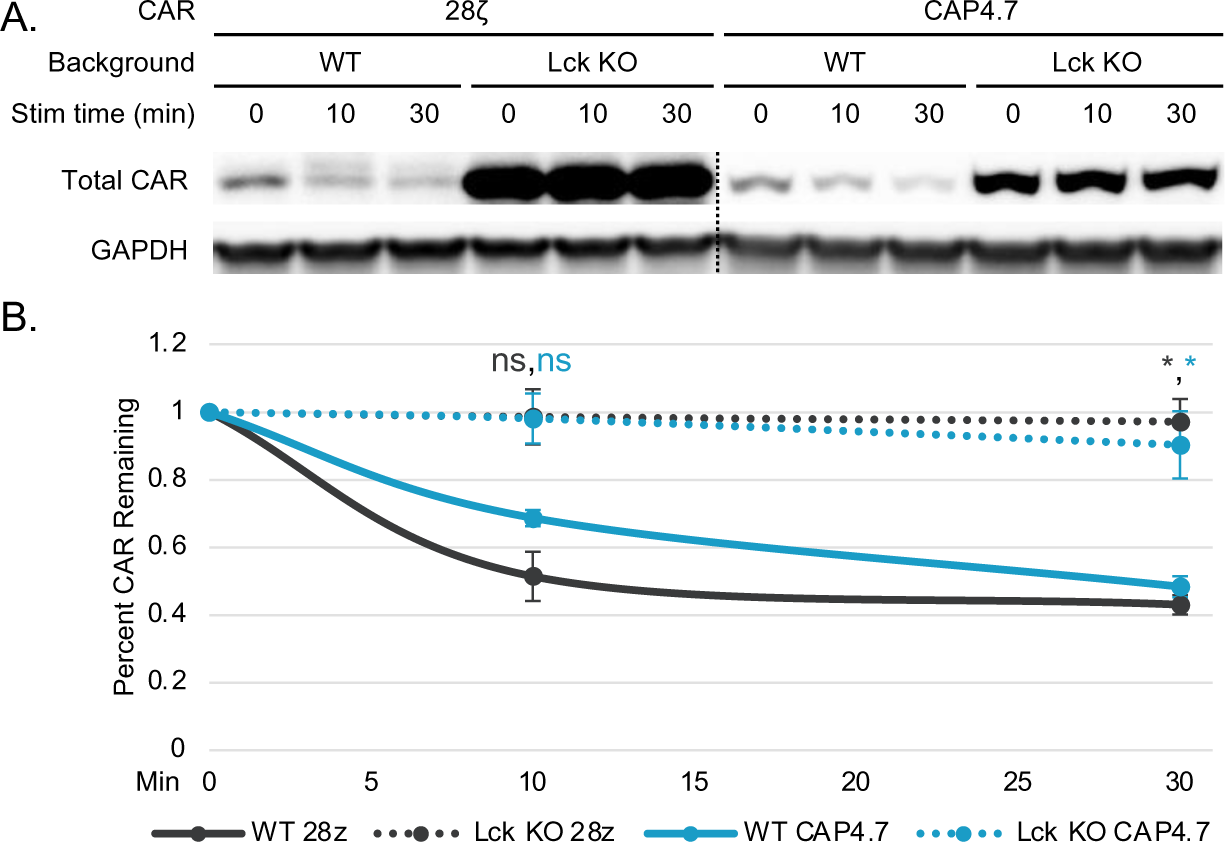
Exploring Drivers of CAR/CAP Degradation. Jurkat-E6.1 cells stably expressing CAR or CAP constructs were mixed with either antigen negative or antigen positive cells. Antigen negative cell mixtures were incubated at 37°C for 10minutes to control for the effect of heating and used as the 0min timepoint. Antigen positive cell mixtures were incubated at 37°C for given amounts of time. Whole cell lysates were then immunoblotted for total CAR. Blot volumes were quantified using Bio-Rad Laboratories’ Image Lab software and normalized to total protein. V = Volume_CAR_/Volume_GAPDH_. Percent CAR Remaining = V/V_maximum_. Data are representative of three independent experiments. **A.** Representative blots of total 28ζ and CAP4.7 levels in WT and Lck KO cells over the signaling time course. **B.** Averaged graphs of phosphorylation curves for different markers of CAR/CAP activation/signaling. Paired Student’s T Tests were performed comparing the total CAR or CAP levels in WT vs Lck KO cells at each timepoint. Bars denote ±SEM. ns: P > 0.05. *: P ≤ 0.05.

## Discussion

CARs have transformed the treatment of blood cancers, but treatment of solid tumors and tumors with low antigen have been less successful due to poor persistence and decreased responsiveness (*5, 33*). A major caveat is the inefficient proximal signaling propagated by CARs (*6, 10, 11*) compared with TCRs. Attempts to improve the signaling properties of CARs by incorporating native TCR elements have shown promise in preclinical models (*34–37*). We focused on incorporating signaling molecules further downstream of the TCR to increase the sensitivity of a CAR molecule and achieve better efficacy. To this end we constructed Chimeric Adapter Proteins (CAPs) that contain the adapters LAT or SLP76 in tandem with the ZAP70 kinase domain. We found that inclusion of adapter proteins caused high antigen-independent activation of T cells, while chimeric molecules containing intracellular ZAP70 domains alone displayed low basal and high antigen-dependent signaling. We proceeded to evaluate these latter ZAP-CARs, labeled CAP4s. While CAP4s appeared similar in *in vitro* assays, second generation CAP4s (CAP4.6 and CAP4.7) outperformed 28ζ-CARs for persistent tumor clearance in an *in vivo* model of leukemia.

CAPs were designed on the rationale that triggering signaling downstream of the TCRζ chain would have the advantage of being more potent because they would bypass the kinetic proofreading steps that define the signaling threshold (McKeithan PNAS 1995) and the inhibitory regulation of upstream molecules subject to negative regulation by inhibitory proteins such as PD1 (*38*). Other recent studies have observed benefits of incorporating downstream TCR signaling components in CAR design (*11, 39*). In the context of CAP, antigen engagement would more directly activate the CAP ZAP70 kinase domain, leading to phosphorylation of critical adapter proteins. These designs represent examples of engineering innovations that have been guided by in-depth biochemical, structural and imaging studies of proximal T cell signaling (*12, 15, 21, 23, 40, 41*). Some of the advantages displayed by CAP4-Ts are their lower tonic signaling than 28ζ-CAR-Ts, reduced T cell differentiation profiles, increased expansion, and finally, a more durable *in vivo* tumor response; these are all properties that suggest significant benefits for translation to clinical settings.

Testing of CAPs and CARs in an *in vivo* model of leukemia revealed striking differences between first and second-generation CAPs. Second-generation CAPs that included a functional CD28 costimulatory domain showed more persistent tumor regression *in vivo,* indicating that the signaling properties conferred by the CD28 costimulatory domain are required for persistent CAP-T cell function. These observations mirror the differences observed between first and second-generation CARs. Both CD28 and 4–1BB costimulatory domains in second generation CAR designs extended T cell survival compared to first generation CARs, but with different characteristics. 28ζ-CARs exhibit higher activity against antigen-low leukemias while BBζ-CAR appear to result in a higher persistence (*26, 42*). In our study second generation CAPs were designed with the 28ζ-CAR backbone, and include the CD28 hinge, TM, and costimulatory domain. Importantly, they show more durable control of tumor progression than the parent CD28ζ-CAR. Thus, signaling properties conferred by ZAP70 domains are responsible for the higher efficacy of second-generation CAPs, with full-length ZAP70 in the CAP4.7 design resulting in the highest T cell expansion and lowest level of terminal differentiation.

Second-generation CAPs showed similar *in vivo* efficacy to 4-1BBζ CAR. While tumor regression and T cell expansion were similar between 4-1BBζ CAR-Ts and second-generation CAP-Ts, surface expression of CAPs was downregulated while that of 4-1BBζ-CAR persisted. This decrease in CAR expression, potentially providing transient rest––a context that has recently been found to restore function in exhausted CAR T cells (*43*), may account for the less differentiated profile of second-generation CAP expressing cells. Incorporation of the 4-1BB domain in CAP designs will allow for a more direct comparison, assessing whether 4-1BB modulates surface CAR expression and more importantly, long-term persistence and function.

Elucidating the signaling properties of chimeric molecules used in immunotherapy is key to understanding differences in the clinical outcomes of patients treated with these different constructs. Several studies have recently been performed to compare CAR and TCR signaling and have shown that CARs have higher and more rapid signaling kinetics compared to TCR signaling (*11, 44–47*), with 28ζ-CAR showing faster activation and a larger magnitude of signaling than 4-1BBζ-CAR (*47*). Using both microscopy and biochemistry we observed that CAPs and downstream signaling proteins exhibit lower levels of phosphorylation but remain activated for longer durations compared with 28ζ -CAR. Thus, CAP signaling more closely resembles the moderate and prolonged signaling that characterizes the TCR. These signaling properties of CAPs provide insights into their enhanced *in vivo* performance and suggest that reduced signal strength and prolonged signaling kinetics are advantageous for chimeric receptor designs.

Though CAR-T therapy has been successful in the clinic, requirements for CAR signaling itself has not been well-defined. To this end we attempted to identify molecules important for CAR and CAP signaling. Both 28ζ -CAR and CAP4s displayed TCR- and Lck-independent signaling. While TCR-independent CAR signaling has been previously reported (*48*), the observation that CAPs, and to a lesser extent 28ζ -CAR, can be triggered without Lck is a novel observation with translational potential. Lck is associated with an exhausted phenotype in 28ζ-CARs, because treating 28ζ-CAR-Ts with the Lck inhibitor Dasatinib, or generating conditions wherein Lck is dephosphorylated, reduce the exhausted phenotype (*43, 49, 50*). Thus, the combination of CAP expression and Lck deletion/inhibition may provide an innovative approach to generate T cells expressing an effective chimeric immunotherapy receptor that undergoes lower levels of degradation and where the resulting T cells exhibit lower terminal differentiation and exhaustion.

In conclusion, our study demonstrates that chimeric receptors that take advantage of signaling molecules in the TCR signaling cascade generate potent and persistent T cell responses against tumors. Incorporation of TCR proximal signaling molecules in CAR designs may provide a new tool in the immunotherapy arsenal.

## Methods

### CAR Lentiviral Vector production and T cell transduction

FMC63-4-1BBζ and FMC63-28ζ−CAR constructs have been previously described. CAP constructs were designed and synthesized followed by cloning into the same lentiviral parent transfer plasmids (pELNS) by VectorBuilder Inc.

CAR or CAP-encoding lentiviral particles were either produced by VectorBuilder Inc or by transient transfection of the Lenti-X 293T lentiviral packaging cell line modified from a previously described method. Briefly, Lenti-X 293T cells were plated into poly-D-lysine-coated 15-cm plates (BD Biosciences). The following day, Lenti-X 293T cells were transfected using Lipofectamine 3000 (Thermo Fisher Scientific) with plasmids encoding the bivalent CAR along with packaging and envelope vectors (pMDLg/pRRE, pMD-2G, and pRSV-Rev). Lentiviral supernatants were harvested at 24 and 48 hr post-transfection, centrifuged at 3,000 rpm for 10 min to remove cell debris, frozen on dry ice, and stored at −80°C.

Human peripheral blood mononuclear cells (PBMCs) from normal donors were obtained with an NIH-approved protocol and activated with CD3 and CD28 microbeads at a ratio of 1:3 (Dynabeads Human T-Expander CD3/CD28, Thermo Fisher Scientific, catalog no. 11141D) in AIM-V media containing 40 IU/mL recombinant IL-2 and 5% FBS for 24 hr. Activated T cells were resuspended at 2 million cells per 2 mL lentiviral supernatant plus 1 mL fresh AIM-V media with 10 μg/mL protamine sulfate and 100 IU/mL IL-2 in 6-well plates. Plates were centrifuged at 1,000 × *g* for 2 hr at 32°C and incubated overnight at 37°C. A second transduction was performed on the following day by repeating the same transduction procedure described earlier. The CD3:CD28 beads were removed on the third day following transduction, and the cells were cultured at 300,000 cells per milliliter in AIM-V medium containing 100 IU/mL IL-2, with fresh IL-2-containing media added every 2–3 days until harvest on day 8 or 9.

### Cytotoxicity Assay

5E4 of target tumor cells in 100 μL RPMI media were loaded into a 96-well plate (Corning BioCoat Poly-L-Lysine 96-Well Clear TC-Treated Flat Bottom Assay Plate). CAR/CAP-T cells were added into the designated well at the indicated ratio. Samples were loaded in triplicate and included T cell-only and tumor-cell-only controls. After 4-6 hr in a 37°C incubator, the plate was scanned for luciferase to monitor cell lysis. The percentage of cell killing at each time point was determined relative to baseline.

### Analysis of Cytokine Production

K562 or K562 CD19 target tumor cells and transduced CAR/CAP^+^ T cells were washed 3 times with PBS and resuspended in RPMI at 1E6 cells per milliliter. 100 μL (1 × 10^5^) tumor cell suspension and 100 μL CAR-T cell suspension was loaded into each well of a 96-well plate with T cell-only and tumor-cell-only controls in triplicates. After 18 hr in a 37°C incubator, a culture supernatant was harvested for detection of the cytokines using ELISA (R&D Systems).

### Flow Cytometry

Cells were washed twice with PBS and stained with either Live/Dead UV L34962or Live/Dead Violet L34955 (Thermo Fisher Scientific) for 30 minutes at 4°C in the dark. Cells were washed in FACS buffer (PBS supplemented with 2% BSA), Fc blocking was performed with a combination of antibodies diluted in FACS buffer with Brilliant Violet Stain Buffer (BD Biosciences) for 30 minutes at 4°C in the dark. Surface expression of CD19-CAR was detected with PE-anti-FMC63 scFv Antibody, Mouse IgG1 (Y45) (Acro Biosystems FM3-HPY53). The following antibodies were used on cells isolated from bleeds and spleen/bone marrow of NSG treated mice: CD3 APC R700 (BD 565119), CD4 BB700 (BD 566392), CD8 APC Cy7 (BioLegend 344714), PD1 PE-Cy7 (BioLegend 329918), Tim3 AF647 (BD 565558), LAG3 BV605 (BioLegend 369324), CD62L BV650 (BD 583808), CD45RA BV786 (BD 565419). Cells were washed in FACS buffer and analyzed by flow cytometry. If cells were not able to be analyzed on the same day, they were fixed in 4% PFA and analyzed by flow cytometry the next day. Flow cytometry was performed on a BD FACS Symphony or BD LSRII and analyzed with FlowJo software version 10.5 or greater (Tree Star).

### In Vivo Studies

All animal procedures reported in this study that were performed by NCI-CCR affiliated staff were approved by the NCI Animal Care and Use Committee (ACUC) and in accordance with federal regulatory requirements and standards. All components of the intramural NIH ACU program are accredited by AAALAC International. Nalm6 cells expressing GFP and luciferase were intravenously (i.v.) injected into NSG mice (NOD.Cg-*PrkdcscidIl2rgtm1Wjl/*SzJ; Jackson Laboratories). Leukemia was detected using the Xenogen IVIS Lumina (Caliper Life Sciences). Mice were injected intraperitoneally with 3 mg D-luciferin (Caliper Life Sciences) and were imaged 4 min later with an exposure time of 30 s for NALM6 and 2 min for PDXs. Living Image Version 4.1 software (Caliper Life Sciences) was used to analyze the bioluminescent signal flux for each mouse as photons per second per square centimeter per steradian.

### Protein Structure Modeling

Refer to Supplemental Methods

### Imaging experiments

Lentiviral constructs utilized include 28ζ CAR-GFP and CAP4.7-GFP and were generated by VectorBuilder. DNA constructs include Zap70-Apple and Grb2-Apple which were described previously (Yi et al., 2019). For the fixed cell imaging experiments, Raji B cells were resuspended in serum-free RPMI and allowed to adhere to a Poly-L-Lysine coated coverslip at 37C for 1 hour. CAR or CAP T cells were pipetted onto them and incubated for 10 minutes. Cells were then fixed with 4% paraformaldehyde for 30 minutes, washed 3x with 1X PBS. Samples were permeabilized in 0.1% Triton-X-100 for 3 min and then incubated in a blocking solution consisting of 10% FBS (Sigma-Aldrich), 0.01% sodium azide (Sigma-Aldrich), and 1 × PBS for 1 h at room temperature (RT). After three washes in 1 × PBS, the cells were stained with primary antibody in blocking solution (anti-SLP-76 pY128 from BD Biosciences, catalog no. 558367 was used at 15μg/ml) for 1 h at RT, followed by secondary antibody in blocking solution (isotype-specific Alexa Fluor-conjugated secondary antibody was used at 1:1000-fold dilution) for 45 min at RT. Samples were then imaged using a spinning disk confocal microscope (Nikon Ti with a Yokogawa CSU-X1 head) operated by the Andor iQ3 software. Acquisitions were performed using a 100× objective (CF1 PlanApo λ 1.45 NA oil), and an EMCCD iXon897 camera (Andor). For immune synapse image analysis, 28ζ-CAR-GFP and CAP4.7-GFP labeled immune synapses in fixed-cell images were manually segmented using the ‘Label’ function in the napari image viewer (*51*). Intensity and volume measurements quantified using a custom Python script using the *scikit-image* library (*52*).

For the live cell TIRF imaging experiments, coverslips were coated with 10ug/ml anti-CD19 antibodies to trigger CAR and CAP T cell activation. Live cell imaging performed as previously described and lag times were calculated as previously described (Yi et al., 2019). The accumulation time’ (*τ_Accum_*) of microcluster intensity *vs* time traces were determined using an in-house written MATLAB (Mathworks Inc.) program. For these, the program utilizes the local slope (by calculating the first derivative) of the background corrected intensity curves. The ‘outset time’ (*τ_Outset_*) is reported as the time corresponding to the first major positive deviation of the local slope above a threshold, estimated from slope fluctuations of pixel intensities without microcluster fluorescence. The ‘accumulation time’ (*τ_Accum_*) was calculated as *τ_Accum_* = *τ_Max_* − *τ_Outset_*, where *τ_Max_* is the time of maximum microcluster intensity.

### Stimulation of CAR expressing Jurkat cell lines

Cultures of CAR expressing Jurkat cell lines were spun down, resuspended in cold RPMI media at 10×10^6^ cells per 100μl, aliquoted into separate Eppendorf tubes for each condition, and put on ice. 5×10^6^ or 10×10^6^ (50 or 100μl of) Jurkat cells were used per condition. Cultures of CD19 negative or positive K562 cell lines were also spun down and resuspended in cold RPMI media at 10×10^6^ cells per 100μl. Equal volumes of K562s were added to each appropriate Jurkat Eppendorf on ice, creating a 1:1 Jurkat-to-K562 ratio. Each sample was then spun down at 300g for 1 minute at 4°C and immediately returned to ice. Each sample was incubated in a 37°C hot water bath for a given amount of time and then immediately lysed on ice.

For the kinetics experiments, two negative control samples were used to control for the effects of incubation: one with Jurkat + K562 CD19^-^ cells lysed immediately post-centrifugation and one with Jurkat + K562 CD19^-^ cells lysed after 10 minutes incubation. For all other stimulation experiments, the negative control samples had Jurkat + K562 CD19^-^ cells lysed after the same incubation time as the experimental samples.

For PP1 inhibition experiments, Jurkats and K562s were pre-treated with 20µM of either DMSO or PP1 for 30 minutes at 37°C and 10^6^ cells per 1ml and returned to ice before stimulating. The media used during stimulations also had either 20µM DMSO or PP1.

For lysis, a 4:1 volume ratio of lysis buffer to media+cells was used, creating whole cell lysate concentrations of 20,000cell/μl (10,000Jurkat/μl + 10,000K562/μl). Samples were then spun down at 4°C at 14,000rpm for 10 minutes, and the supernatants were collected for use as WCLs. Lysis buffer recipe: 25mM TRIS pH 8.0, 150mM NaCl, 1% NP-40, 5mM EDTA, 1mM Na_3_VO_4_, 1X cOmplete^™^ (Roche Cat no.: 11836153001).

### Immunoblotting under Reducing and Non-Reducing conditions

For regular blots, a 4:1 ratio of WCL to 5x sample buffer was used to prepare samples for blotting. 5x reducing sample buffer recipe: 50mM TRIS pH 8.0, 5mM EDTA, 5% SDS, 50% glycerol, 5mM Na_3_VO_4_, 0.05% bromophenol blue, 50mM DTT, 850mM βME. For non-reducing blots, the same protocol and reagents were used except the sample buffer did not contain DTT or βME.

After mixing with sample buffer, samples were heated at 95°C for 5 min. 2.6×10^5^ cell equivalents were separated by SDS/PAGE using 10% Criterion Precast polyacrylamide gels. The separated proteins were then transferred to nitrocellulose membrane and the membrane was blocked for 1 hour at room temperature using TBST [10 mM Tris (pH 8.0), 150 mM NaCl, and 0.05% Tween 20] with 5% milk and 1% BSA. The membranes were incubated overnight at 4 °C with primary Abs diluted in TBST with 5% milk, 1% BSA, followed by a 60-min incubation at room temperature with the appropriate secondary Ab diluted in TBST with 5% milk, 1% BSA. The blots were then visualized by chemiluminescence and quantified using Bio-Rad’s Image Lab software.

Antibodies used for immunoblotting: pCAR (28ζ): BD Biosciences 558402; pCAP (CAP4.6 + CAP4.7): Cell Signaling 2701; Total CAR (28ζ): Santa Cruz Biotechnology sc-1239; Total CAR (CAP4.6 + CAP4.7): Abcam ab32429; pLck: Cell Signaling 2101; pZeta: BD Biosciences 558402; pZAP: Cell Signaling 2701; pLAT: BD Biosciences 558363; pSLP76: BD Biosciences 558367; pPLCγ1: Cell Signaling 2821; pERK: Cell Signaling 4370; pAKT: Cell Signaling 4060; GAPDH: Cell Signaling 2118

### Quantification of western blots and calculation of protein phosphorylation and dephosphorylation rates

The background corrected and normalized Intensity *vs* Time data from individual Western blot gel runs were fitted to an expression modeling exponential rise and decay, given by,

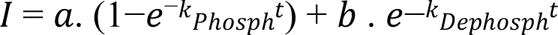

where *k_Phosph_* and *k_Dephosph_* are the respective phosphorylation and dephosphorylation rates, *t* is the time and *a*,*b* are amplitude coefficients. The fittings were done using a non-linear regression routine implemented in MATLAB. Mean values of the phosphorylation and dephosphorylation rates for each protein were calculated from repeat experiments and plotted as bar graphs, along with their corresponding standard deviation.

## ACKNOWLEDGEMENTS

This research was supported by the Intramural Research Program of the NIH, NCI, CCR. We thank the CCR/LGI Flow Cytometry Core for flow analyzers. We thank Dr. Jiyao Wang from NCBI for his help in providing iCn3D links improvements.

## AUTHOR CONTRIBUTIONS

L.B., T.M., H.Q., N.A., J.Y. and K.M. performed the experiments; L.B., J.Y., N.T. and L.E.S. designed the study; L.B., N.A., S.P. and A.T. performed image analysis; L.B., H.Q., M.L. and H.Y. performed flow analyses; P.Y. and R.C. performed structural studies; L.B., T.M. and R.C. prepared figures, L.B. wrote the manuscript with comments from J.Y., H.Y., N.T. and L.E.S.

---

**Figure S1.**
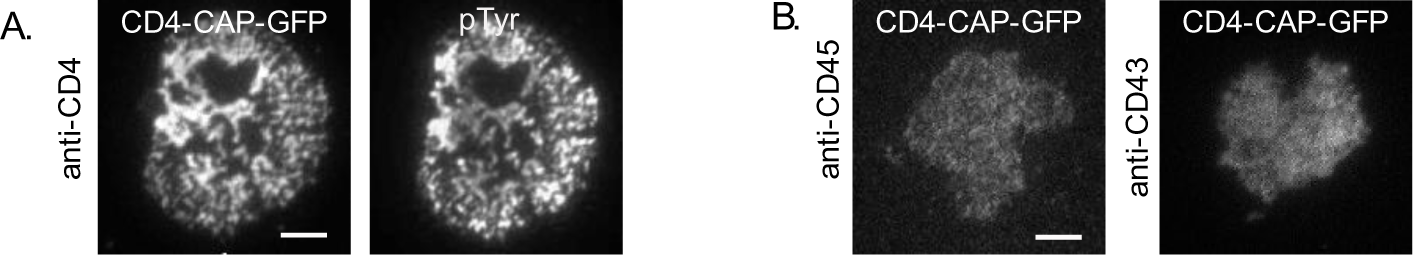
Chimeric Adapter Proteins (CAPs) cluster specifically upon antibody binding of extracellular domain. **A and B.** TIRF images of microclusters formed in Jurkat T cells expressing CD4-CAP-GFP and activated on coverslips coated with indicated antibodies. In **A**, cells were fixed and immunostained with phosphotyrosine antibody to detect activated microclusters. Scale bars in images, 2 μm.

**Figure S2.**
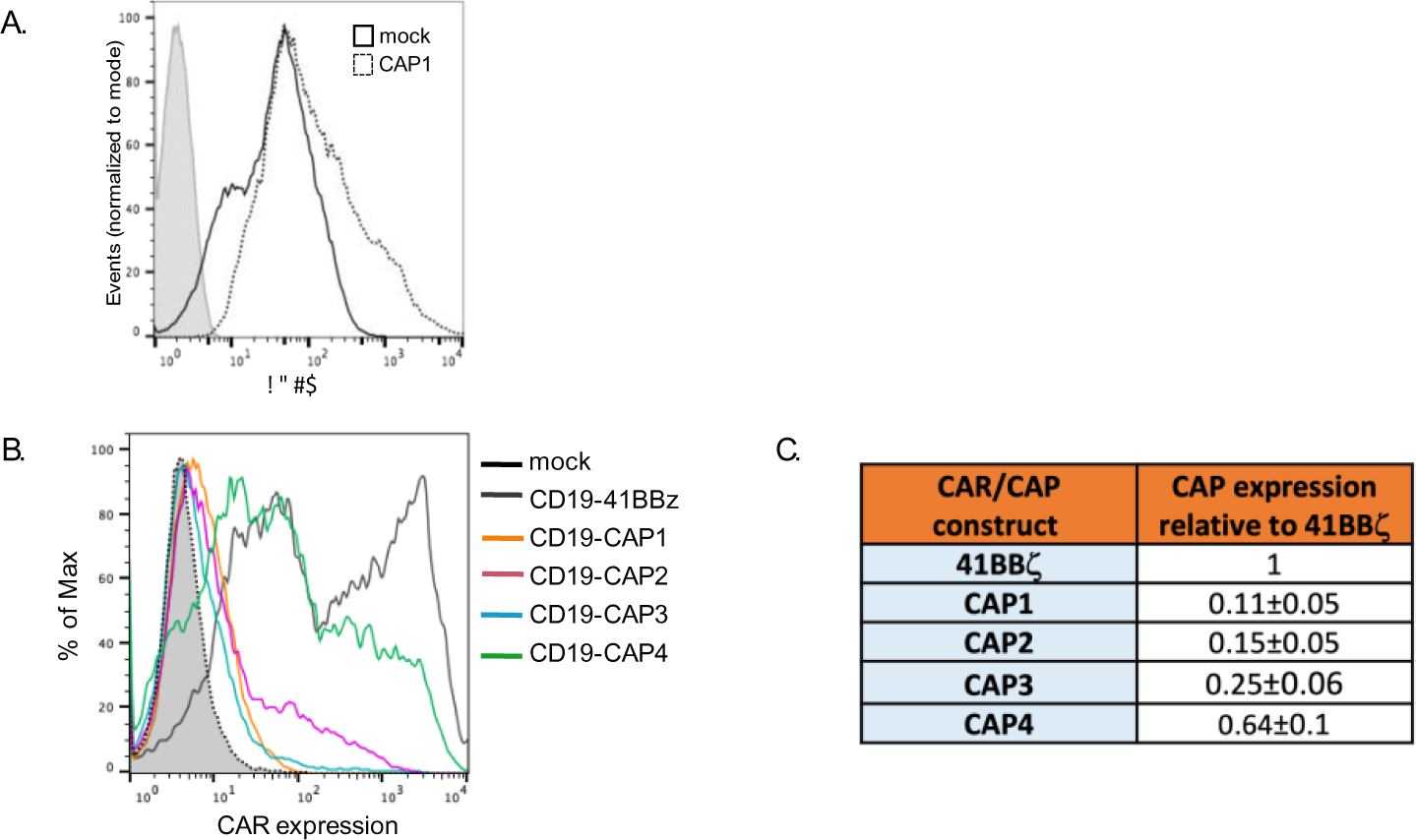
Surface expression of molecules in T cells expressing CAPs. **A.** Basal CD69 expression in mock transduced Jurkat cells and Jurkat cells transduced with CD19-CAP1. **B.** Representative histograms of CD19 scFv surface expression in PBMCs transduced with the indicated CD19-41BBζ CAR or CAP constructs. **C.** T cells were transduced with the different CAP constructs and the surface expression of each CAP relative to 4-1BBζ CAR in 3 different donors are presented as means±SD.

**Figure S3.**
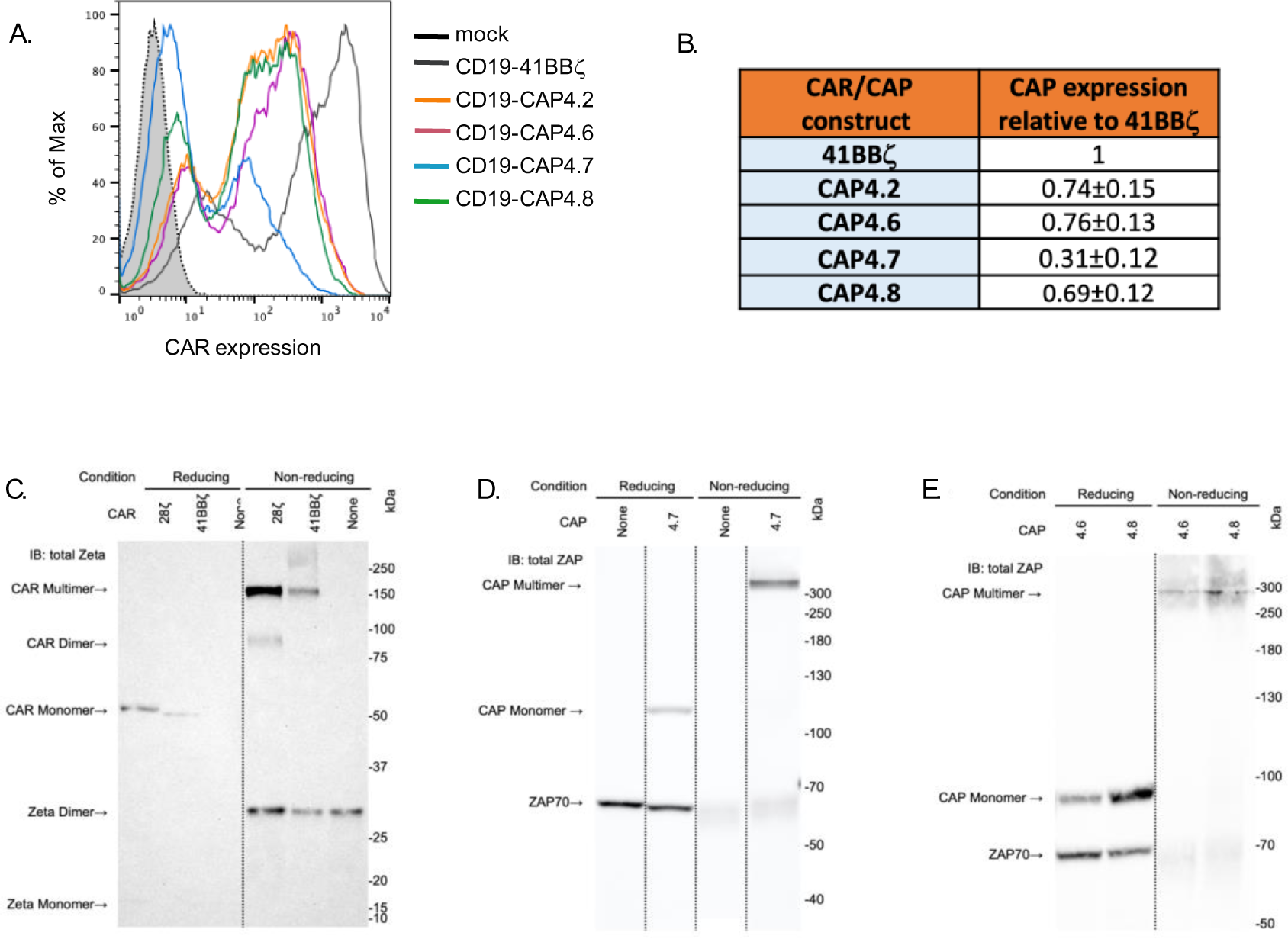
Surface expression of CAP4s and western blots to detect CARs and CAPs under reducing and non-reducing conditions. **A.** Representative histograms of cell surface CD19 scFv expression in T cells transduced with the indicated CD19-CAR and CAP4 constructs. **B.** T cells were transduced with the different CAP constructs and the surface expression of each CAP relative to 4-1BBζ CAR in 3 different donors are presented as means±SD. **C-E.** Western blots to detect CARs and CAPs. Jurkat E6.1 cells that were either untranduced or stably expressing CAR or CAP constructs were lysed and then immunoblotted under reducing or non-reducing conditions. CARs were detectable with anti-Zeta antibody and CAPs were detectable with anti-ZAP antibody. **C.** Total ζ blot of CAR and control cells. Endogenous TCRζ detectable at ∼15kDa under reducing conditions and ∼29kDa under non-reducing conditions. 28ζ CAR detectable at ∼52kDa under reducing conditions and ∼92kDa and ∼170kDa under non-reducing conditions. 41BBζ CAR detectable at ∼51kDa under reducing conditions and ∼164kDa under non-reducing conditions. **D.** Total ZAP blot of CAP and control cells. Endogenous ZAP detectable at ∼68kDa under reducing conditions and ∼62kDa under non-reducing conditions. CAP4.7 detectable at ∼117kDa under reducing conditions and above 300kDa marker under non-reducing conditions. **E.** Total ZAP blot of CAP cells. Endogenous ZAP detectable at ∼68kDa under reducing conditions and ∼66kDa under non-reducing conditions. CAP4.6 detectable at ∼90kDa under reducing conditions and ∼292kDa under non-reducing conditions. CAP4.8 detectable at ∼91kDa under reducing conditions and ∼292kDa under non-reducing conditions.

**Figure S4.**
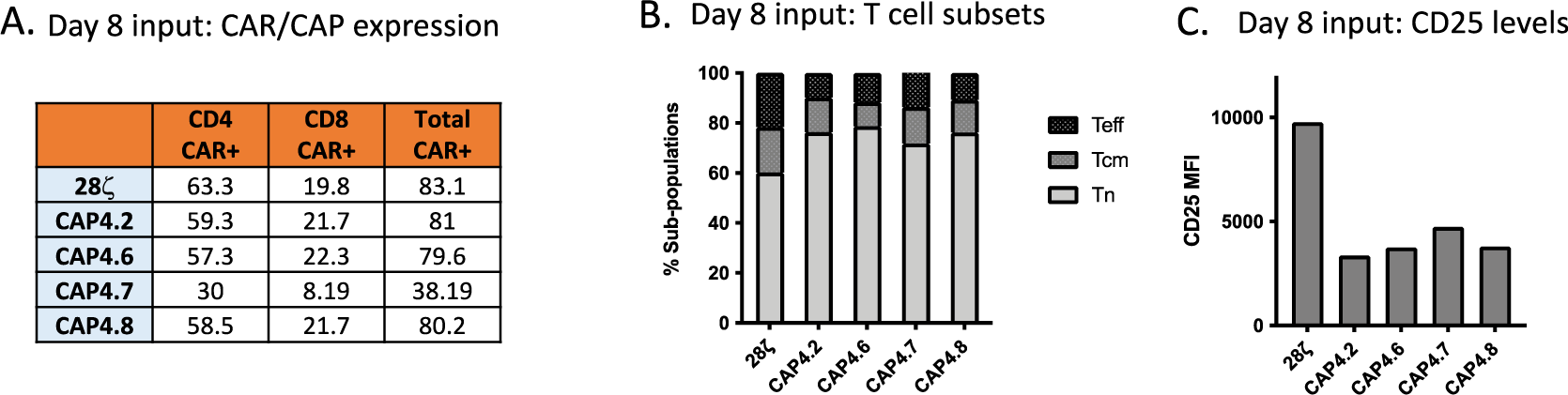
Analysis of transduced PBMCs on Day8 used in *in vivo* experiment in Figure 4. **A.** Table showing the percentages of CAR^+^ or CAP^+^ T cells within the CD4 and CD8 subsets. **B.** Flow cytometric evaluation of CD25 surface levels in CAR^+^ or CAP^+^ T cells. **C.** The percentages of naïve (T_n_), central memory (T_CM_), and effector T cells (T_eff_) within the CAR^+^ or CAP^+^ T cell subset was evaluated by flow cytometry as follows: T_n_ (CD62L^+^CD45RA^+^), T_cm_ (CD62L^+^, CD45RA^-^), T_eff_ (CD62L^-^).

**Figure S5.**
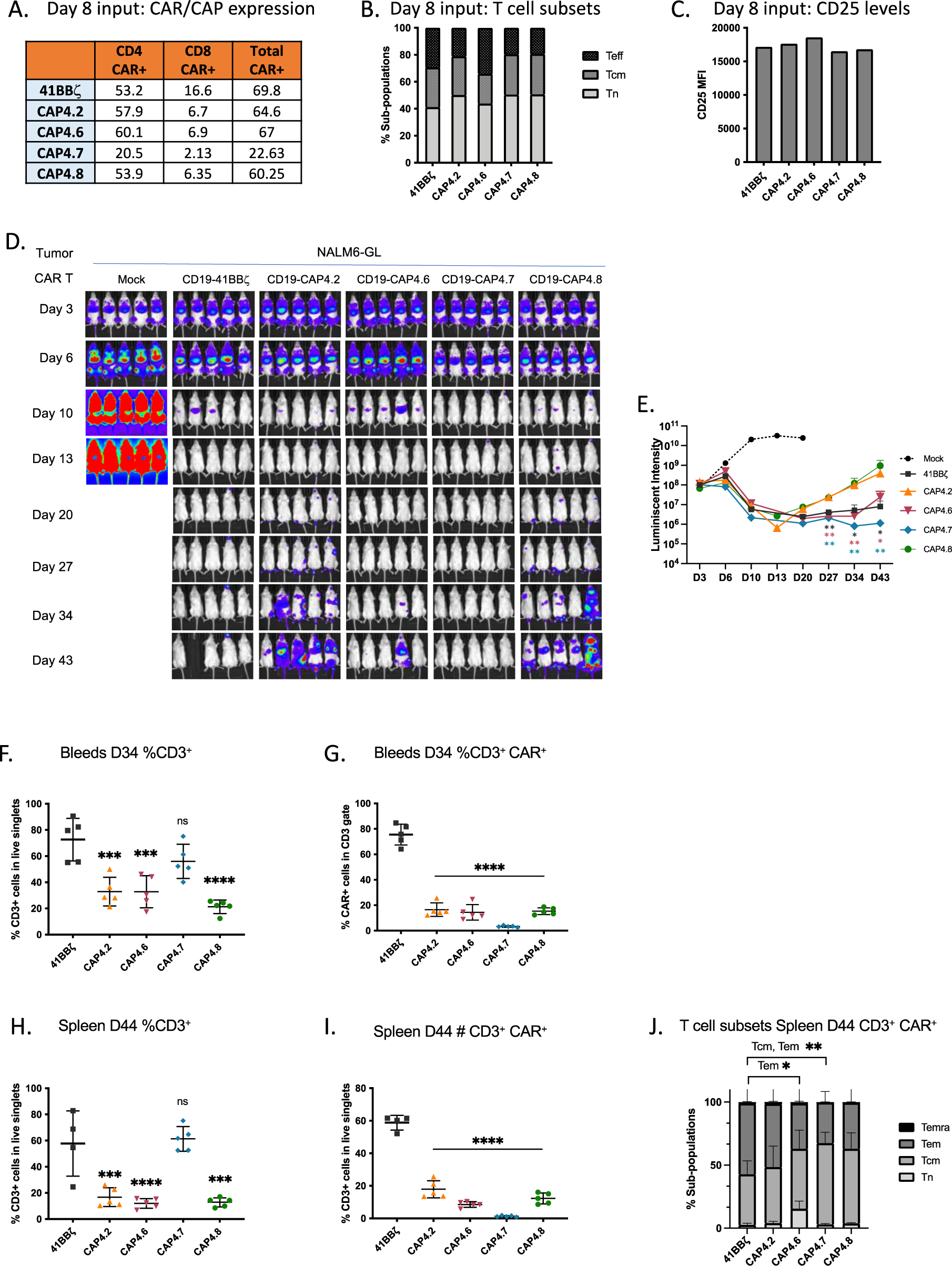
Comparison of CD19-CAP4 constructs with 41BBz-CAR in an *in vivo* NSG leukemia model. **A.** Table showing the percentages of CAR^+^ or CAP^+^ populations in CD4 and CD8 subsets. **B.** Flow cytometric evaluation of CD25 surface levels in in CAR^+^ or CAP^+^ T cells. **C.** The percentages of naïve (T_n_), central memory (T_CM_), and effector T cells (Teff) within the CAR^+^ or CAP^+^ T cell subset was evaluated by flow cytometry as follows: T_n_ (CD62L^+^CD45RA^+^), T_cm_ (CD62L^+^, CD45RA^-^), T_eff_ (CD62L^-^). **D.** NSG mice were engrafted with NALM6 leukemia cells as in Figure 4A/4B, and at day 3, mice were adoptively transferred with T cells transduced with the control CD19-41BB vector or CAP4.2, 4.6, 4.7, or 4.8 constructs (3E6). Leukemia growth was evaluated at the indicated timepoints by bioluminescent imaging (BLI) and IVIS images images are shown. **E.** Quantification of the BLI radiance data for each individual mouse is presented. Bars denote ±SEM. Statistical differences were assessed using a Mann Whitney t-test comparing 4-1BBz-CAR, CAP4.6 or CAP4.7 with CAP4.2. *: P ≤ 0.05; **: P ≤ 0.01. **F and G.** Flow cytometric analysis of peripheral blood on Day 34. **H-J.** Flow cytometric analysis of Spleen on Day 44. T cell subsets were evaluated as follows: T_n_ (CD62L^+^CD45RA^+^), T_cm_ (CD62L^+^, CD45RA^-^), T_em_ (CD62L^-^CD45RA^-^) and T_emra_ (CD62L^-^CD45RA^-^) Two way Anova analysis was performed. Bars denote ±SEM. ns: P > 0.05; *: P ≤ 0.05; **: P ≤ 0.01; ***: P ≤ 0.001. ****: P ≤ 0.0001.

**Figure S6.**
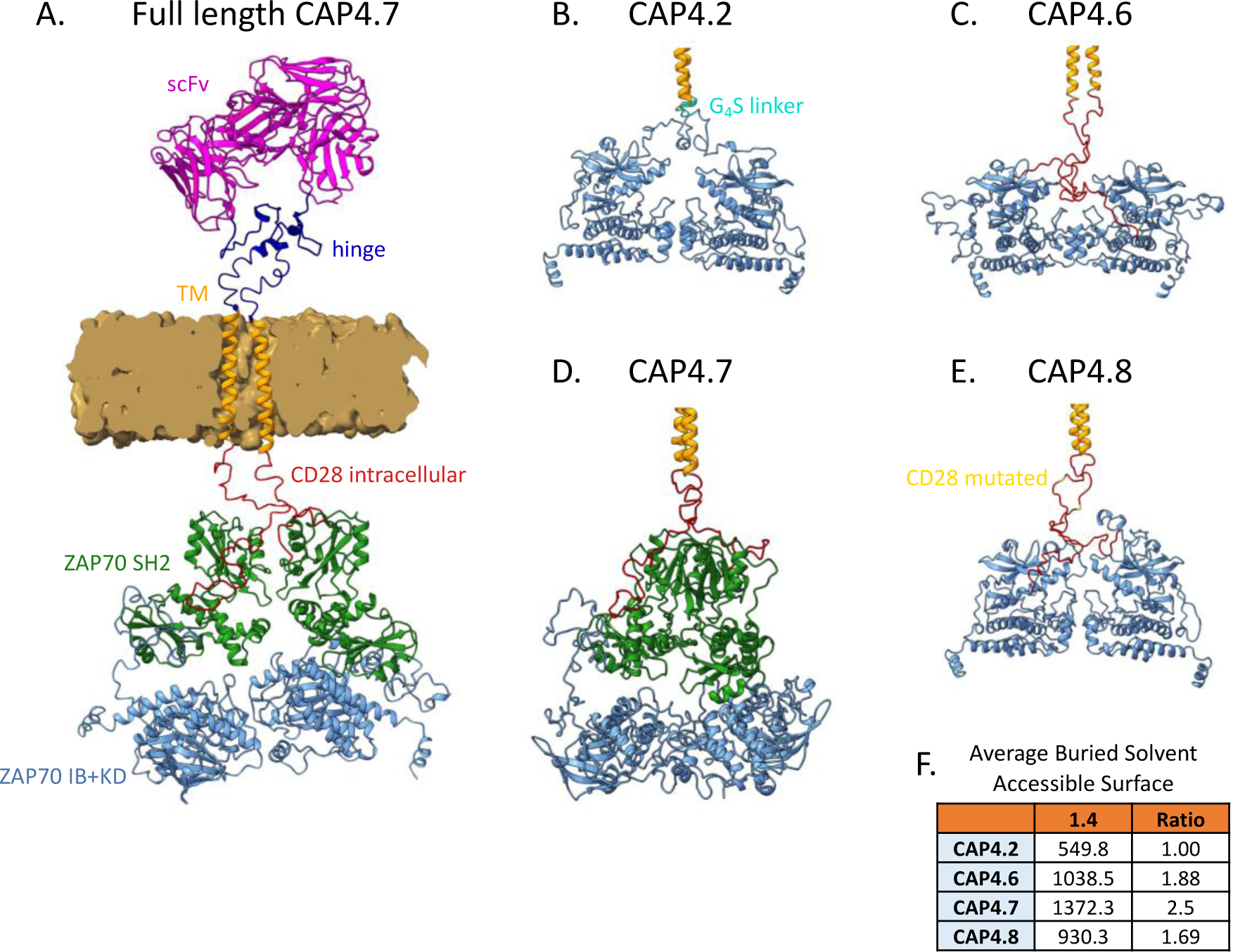
Structural Models of CAP4 constructs. **A.** Full length model of CAP4.7 in the plasma membrane (brown). scFv in pink; hinge in dark blue; transmembrane domain (TM) in yellow; CD28 intracellular domain in red; ZAP70 SH2 domains in green; ZAP70 IB+KD domain in pale blue. **B-D.** Intracellular regions of CAP4.2 **(B)**, CAP4.6 **(C)**, CAP4.7 rotated 90° clockwise along the vertical axis compared to A **(D)**, and CAP4.8 **(E)** are shown. **F.** Table of Average Buried Solvent Accessible Surface numbers and ratios (BSAS) for CAP4 series constructs.

**Figure S7.**
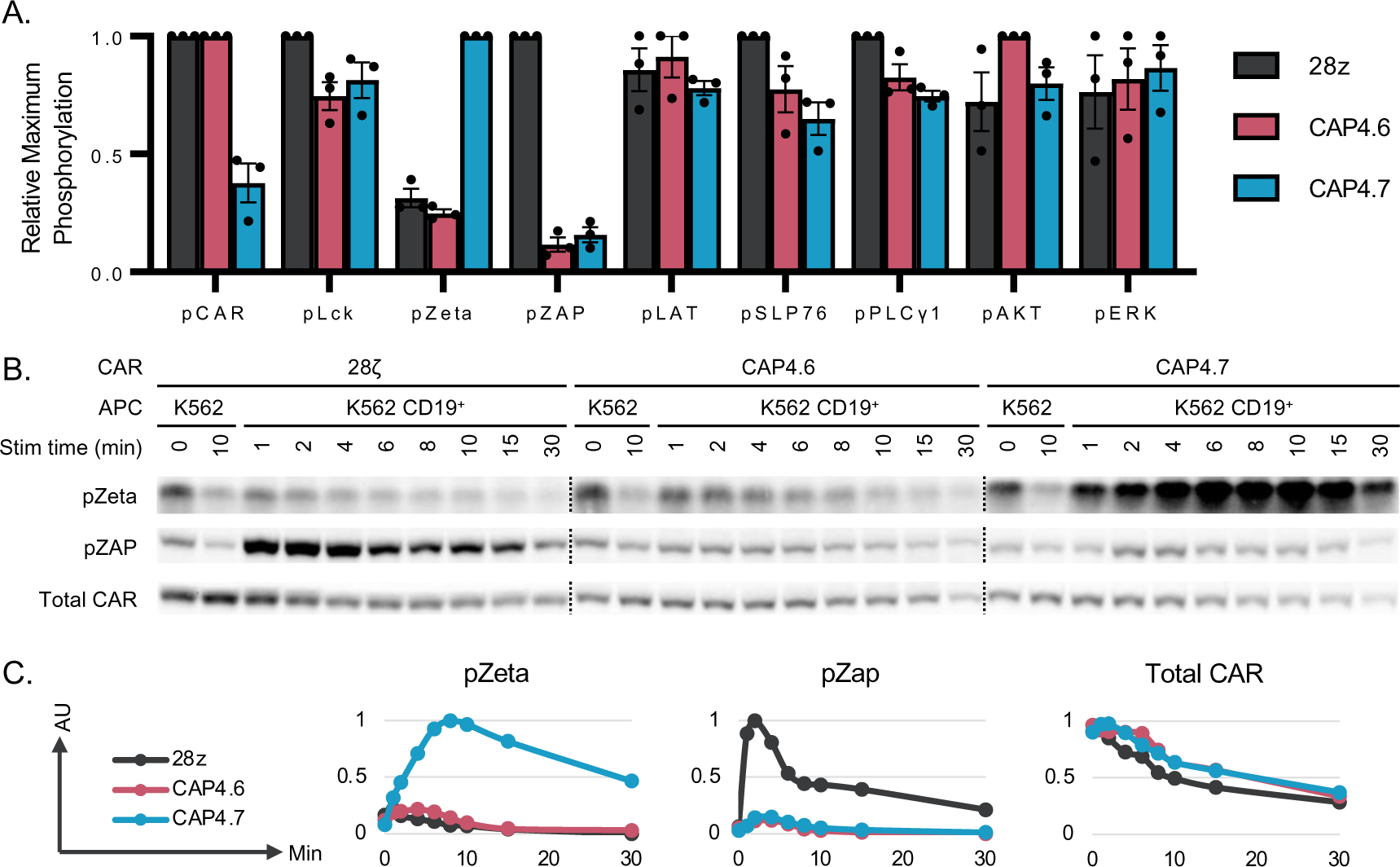
**A.** Averaged maximum phosphorylation intensities of each marker for each construct. **B.** Representative blots of 28ζ, CAP4.6, and CAP4.7 signaling kinetics and total CAR levels. **C.** Averaged graphs of phosphorylation curves for different markers of CAR/CAP signaling. For Total CAR graph, arbitrary units show percent CAR remaining: AU = V/V_maximum_.

**Figure S8.**
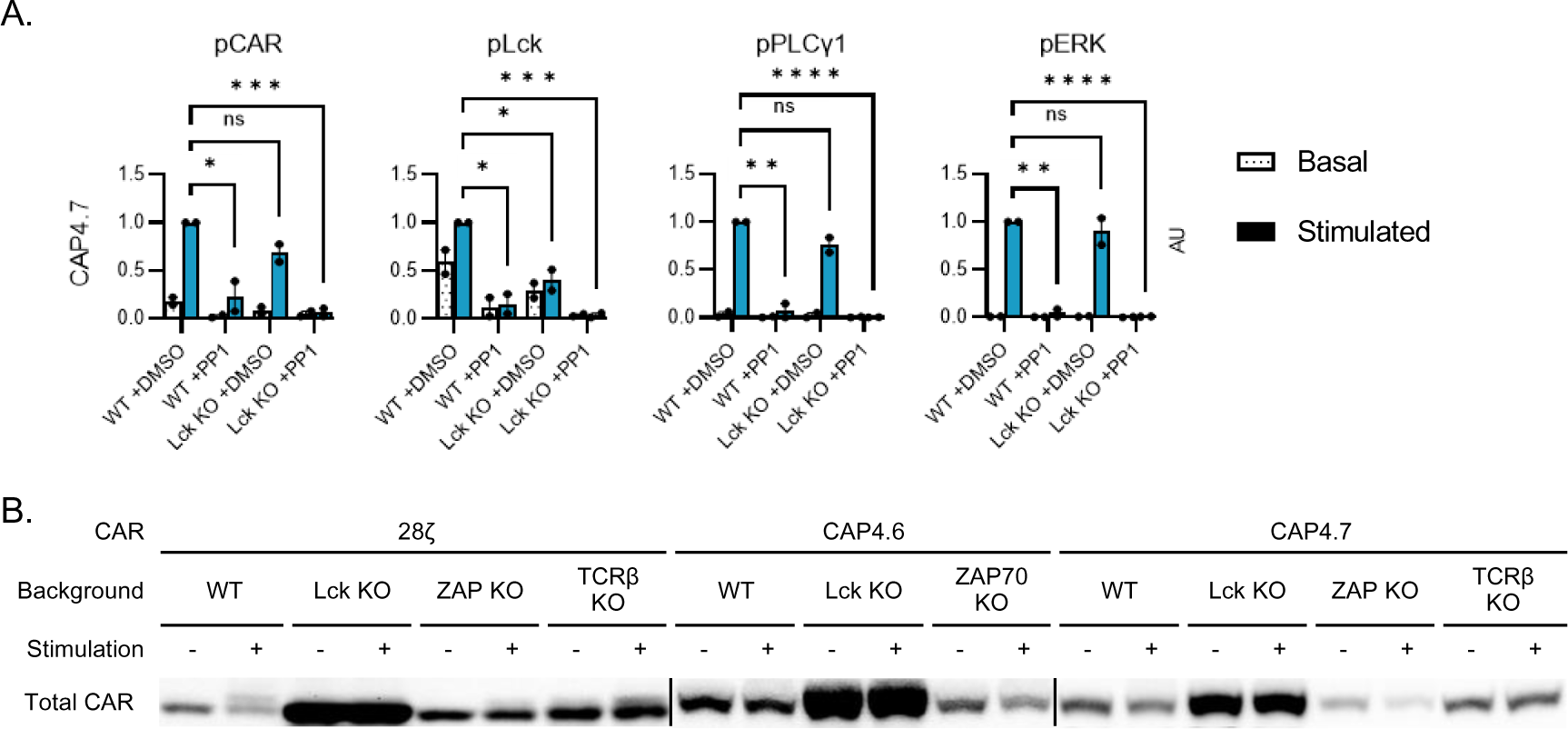
**A.** Averaged graphs of phosphorylation intensity for different markers of CAP4.7 activation/signaling in the presence or absence of Lck and incubation with either DMSO or PP1. Homoscedastic Student’s T Tests were performed comparing the phosphorylation levels of activation/signaling markers in stimulated WT vs Lck KO and DMSO-treated vs PP1-treated samples. Data are representative of two independent experiments (N=2). Bars denote ±SEM. ns: P > 0.05; *: P ≤ 0.05; **: P ≤ 0.01; ***: P ≤0.1 ****: P ≤ 0.0001. **B.** Blot of total 28ζ, CAP4.6, and CAP4.7 levels in WT, Lck KO, ZAP KO, and TCR KO cell lines. Blot is representative of three independent experiments.

## Supplemental Methods

### Molecular Modeling

CAR models were generated following a prescription we previously used to simulate CARs [1]. The procedure relies on the combination of multiple modeling methods, including I-TASSER [2], Rosetta [3], Phyre2 [4], Swiss-Model [5], YASARA [6], and AlphaFold2 [7]. Finally, the structures of all segments (i.e., intracellular, trans-membrane -TM-, extracellular) obtained from all modeling engines were combined using YASARA v20.4.24 HM_build macro to obtain models with better statistics. The hybrid section models were further refined against all templates using FRMD (Feedback Restrain Molecular Dynamics) [8,9] to improve Z-scores and then manually combined and inserted in a model membrane. The final models were tested using MD trajectories. These calculations were performed using YASARA with the Amber14 force field, including the Lipid14 set and standard parameters used in the md_membrane macro. Simulations were performed by inserting the models in membranes made of 20% cholesterol and 80% phospholipids (33% phosphatidylethanolamine, 33% phosphatidylcholine, and 14% phosphatidylserine) following the suggested values [6], representing the behavior of cytoplasmatic membranes. This work was developed concurrently with the advent of AlphaFold2[7], which was the only modeling tool tested for producing stable dimers for the intracellular domains.

Some significant differences were observed in the behavior of the models when comparing the CAP4 group, showing a stable behavior (Cα <RMSF> < 3Å for for trajectories over 50 ns), while 28ζ and 4-1BBζ models were not stable. Attempts at manually stabilizing 28ζ and 4-1BBζ models proved unfruitful. Further analysis of the unfolding of the frustrated models led to the observation of a rapid deterioration of the TM motif during the unwinding of the models’ intracellular domains. This behavior was further explored by performing equilibrium molecular dynamics calculations of the TM itself and then challenging the model using steered dynamics (Yasara md_runsteered) [6]. The stabilization trajectory of the TM domain was conducted for 50 ns, followed by a 1000 ns run using the YASARA MD_run macro in the NPT ensemble. Model stability was assessed using the YASARA MD_analyze macro. The equations of motion for all steered dynamics simulations were integrated with multiple timesteps of 1.25 fs for bonded interactions and 2.5 fs for non-bonded interactions at a temperature of 298K and a pressure of 1 atm (NPT ensemble) using algorithms described in detail previously [10]: after an equilibration time of 3 ps, a minimum acceleration of 2000 pm/ps2 was applied to all steered atoms together with the non-bonded forces (every 2.5 fs).

Considering the steered mass (M) in Dalton, this results in a pulling force of [2000*M*0.00166] picoNewton. The pulling direction was defined by a vector manually provided to point in a previously defined direction. The standard procedure call for a maximum distance (displacement) to be continuously updated. If it did not increase for 400 simulation steps (i.e., the pulling got stuck), the acceleration was increased by 500 pm/ps2. As soon as the maximum distance grew faster than MaxDisSpeed =4000 m/s (i.e., a barrier was overcome), the acceleration is scaled down by a factor of 1-(1-4000/MaxDisSpeed)^2^, but not below the initial minimum. This check is done every 20 simulation steps. Barriers can be qualitatively estimated during this process. To our surprise, the TM model rapidly unwinds when challenged from the intracellular side using the standard steering protocol and pulling K316 towards the inside in a perpendicular direction to the membrane surface. We modified the default protocol to limit the pulling acceleration to 1000 pm/ps2, yet the TM model still losses 25% of its helical content in 6.5 ns. Conversely, if the same protocol is applied to the extracellular side of the TM domain (pulling F287 away from the membrane pointing toward the outside), the TM domain remains more stable, retaining >90% of its helical content (>15ns trajectory). These results can be compared to the TM equilibrium trajectory, which is stable over a 1-microsecond trajectory (average helical content > 95%). A more systematic modeling analysis of different TM domain properties is outside this presentation’s scope and will be presented elsewhere.

However, modeling results should be considered cautiously, especially when experimental structural verification is unavailable. Nevertheless, when taken together, these early observations suggest a larger possible impact of the intracellular domain stability on the stability of the CAR overall. A more detailed exploration of the MD stable CAP4 series AlphaFold2 models further supports this observation. The expected stability of the CAP4 intracellular dimer models can be estimated from the AlphaFold2 dimer averaged buried solvent accessible surface (BSAS), which quantifies the surface area that is accessible by solvent in a monomer and becomes buried with the formation of the dimer. It can be used as a rough estimation of the complex formation stability. [BSAS values were estimated using Chimera 1.16 [11] using default parameters. CAP4.2 shows the smallest BSAS (549.8 Å^2^), followed by CAP4.8 (930.3 Å^2^), CAP4.6 (1038.5 Å^2^) and CAP4.7 (1372.3 Å^2^)]. CAP4.7 AlphaFold2 intracellular dimer arrangement reveals a significant domain rearrangement through the interdomain linker between the C-SH2 domain and the kinase domain.

